# Lsm12 is an NAADP receptor and a two-pore channel regulatory protein required for calcium mobilization from acidic organelles

**DOI:** 10.1101/2020.05.21.109850

**Authors:** Jiyuan Zhang, Xin Guan, Jiusheng Yan

## Abstract

Nicotinic acid adenine dinucleotide phosphate (NAADP) is a potent Ca^2+^-mobilizing second messenger which uniquely mobilizes Ca^2+^ from acidic endolysosomal organelles. However, the molecular identity of the NAADP receptor remains unknown. Given the necessity of the endolysosomal two-pore channel (TPC1 or TPC2) in NAADP signaling, we performed affinity purification and quantitative proteomic analysis of the interacting proteins of NAADP and TPCs. We identified an Sm-like protein Lsm12 complexed with NAADP, TPC1, and TPC2. Lsm12 directly binds to NAADP via its Lsm domain, whereas TPC-containing membranes and isolated TPCs lose their affinities to NAADP in the absence of Lsm12. Lsm12 is essential and directly involved in NAADP-evoked TPC2 activation and Ca^2+^ mobilization. These findings reveal a putative RNA-binding protein to function as an NAADP receptor and a TPC regulatory protein and provides a molecular basis for understanding the mechanisms of NAADP signaling.

## INTRODUCTION

Intracellular Ca^2+^ signaling, which occurs via changes or oscillation in cytosolic Ca^2+^ concentration, controls almost every aspect of cellular functions and physiological processes. Ca^2+^ mobilization from intracellular stores mediated by second messengers plays a critical role in the regulation of cytosolic Ca^2+^ levels. Among the three known Ca^2+^-mobilizing second messengers in mammalian cells, nicotinic acid adenine dinucleotide phosphate (NAADP), which differs from the enzyme cofactor nicotinamide adenine dinucleotide phosphate (NADP) by a hydroxyl group (**Fig. S1**), is the most potent as it is effective in the low nanomolar range ^1-3^. The other two Ca^2+^-mobilizing second messenger molecules, inositol 1,4,5-trisphosphate (IP_3_) and cyclic ADP-ribose (cADPR), are known to mobilize Ca^2+^ from the endoplasmic reticulum (ER) Ca^2+^ stores by activating IP3 receptors and ryanodine receptors, respectively. In contrast, NAADP mobilizes Ca^2+^ from acidic organelles of endosomes and lysosomes (endolysosomes) through an as yet poorly understood mechanism ^4^ (**Fig. S1**).

NAADP signaling is broadly present in different mammalian cells, involved in many cellular and physiologic processes, and implicated in many diseases including lysosomal storage diseases, diabetes, autism, and cardiovascular, blood, and muscle diseases ^4,5^. Accumulating evidence indicates that endolysosomal two-pore channels (TPCs) are necessary for NAADP-evoked Ca^2+^ release. TPCs are homodimeric cation channels consisting of 2 functional isoforms, TPC1 and TPC2, in humans and mice. Manipulation of TPC expression in cell lines results in changes in NAADP-evoked Ca^2+^ release ^6-9^. Cells’ response to NAADP is eliminated in TPC knockout mice ^6,10,11^ and rescued by re-expression of TPCs in TPC1/2 knockout mice ^11^. NAADP sensitivity and Ca^2+^ permeability were reported for TPC currents recorded on isolated enlarged endolysosomes ^12,13^ and reconstituted lipid bilayers ^14,15^. Thus, TPCs had been considered the most likely candidates of NAADP-responsive Ca^2+^ release channels on endolysosomal membranes. However, other researchers reported that TPCs were not only fully insensitive to NAADP but also Na^+^-selective channels with very limited Ca^2+^ permeability in conventional patch-clamp recordings of exogenous mammalian TPCs expressed on whole enlarged endolysosomes ^16^, plasma membranes ^16,17^, or plant vacuoles ^18,19^. Furthermore, TPCs were not labeled by a photoreactive NAADP-analogue ^20^. Thus, the molecular identity of the NAADP receptor remains elusive and the molecular basis of NAADP-evoked Ca^2+^ release remains controversial and poorly understood.

## RESULTS

### Identification of Lsm12 as an Interacting Partner of both NAADP and TPCs

Given the essential role of TPCs in NAADP signaling, we hypothesize that the NAADP receptor is part of a TPC-containing NAADP signaling multiprotein complex (**Fig. S1**) in which the receptor interacts with the channels either directly as an accessory protein as previously proposed ^21,22^ or indirectly via a different mechanism. We chose the human embryonic kidney (HEK) 293 cell line as a mammalian cell model for identification and functional characterization of the NAADP receptor because of its low endogenous TPC1 and TPC2 expression ^6^ and robust NAADP-evoked Ca^2+^ release upon the heterologous expression of TPCs ^6-9^. To identify the most likely protein candidate of the NAADP receptor, we used TPC1, TPC2, and NAADP as baits in affinity purification/precipitation and a stable isotope labeling by amino acids in cell cultures (SILAC)-based quantitative proteomic approach to identify their mutually interacting proteins (**Fig. 1a**). To immobilize NAADP for affinity precipitation of its interacting proteins, we crosslinked NAADP to adipic acid dihydrazide-agarose via pyridine ribose, as described previously ^23^ (**Fig. 1b**). Upon transient expression of human TPC1 and TPC2 in HEK293 cells, we used immobilized NAADP to pull down the NAADP-interacting proteins and an anti-FLAG antibody to pull down the FLAG-tagged TPC1 and TPC2 and their interacting proteins (**Fig. 1a**). Putative interacting proteins were identified by liquid chromatography-tandem mass spectrometry (LC-MS/MS) as differential proteins in the test and control samples that were differentially labeled by heavy (^13^C) and light (^12^C) isotopes in Arg and Lys (**Fig. 1a)**. Among the four lists of identified putative interacting proteins, an Sm-like (Lsm) protein, Lsm12, was uniquely positioned as an interacting protein shared by TPC1, TPC2, and NAADP (both TPC1- and TPC2-expressing cells) (**Fig. 1c and Table S1**). As demonstrated by the large MS peak ratios (≥3) of heavy (^13^C6-Arg/Lys) and light (^12^C6-Arg/Lys) labeled peptides (**Fig. 1d and Fig. S2)**, Lsm12 was differentially pulled down in the test and in the negative control sample preparations in all four conditions (protein samples of TPC1-eGFP-FLAG or TPC2-eGFP-FLAG expressing cells pulled down by immobilized NAADP and by an anti-FLAG antibody). Immunoblotting confirmed the presence of Lsm12 in the TPC-interactomes, that were pulled down by an anti-FLAG antibody from TPC1-FLAG and TPC2-FLAG expressing cells (**Fig. 1e**); and in the NAADP-interactomes, that were pulled down by immobilized NAADP from both TPC1- and TPC2-expressing cells (**Fig. 1f**).

**Figure 1.**
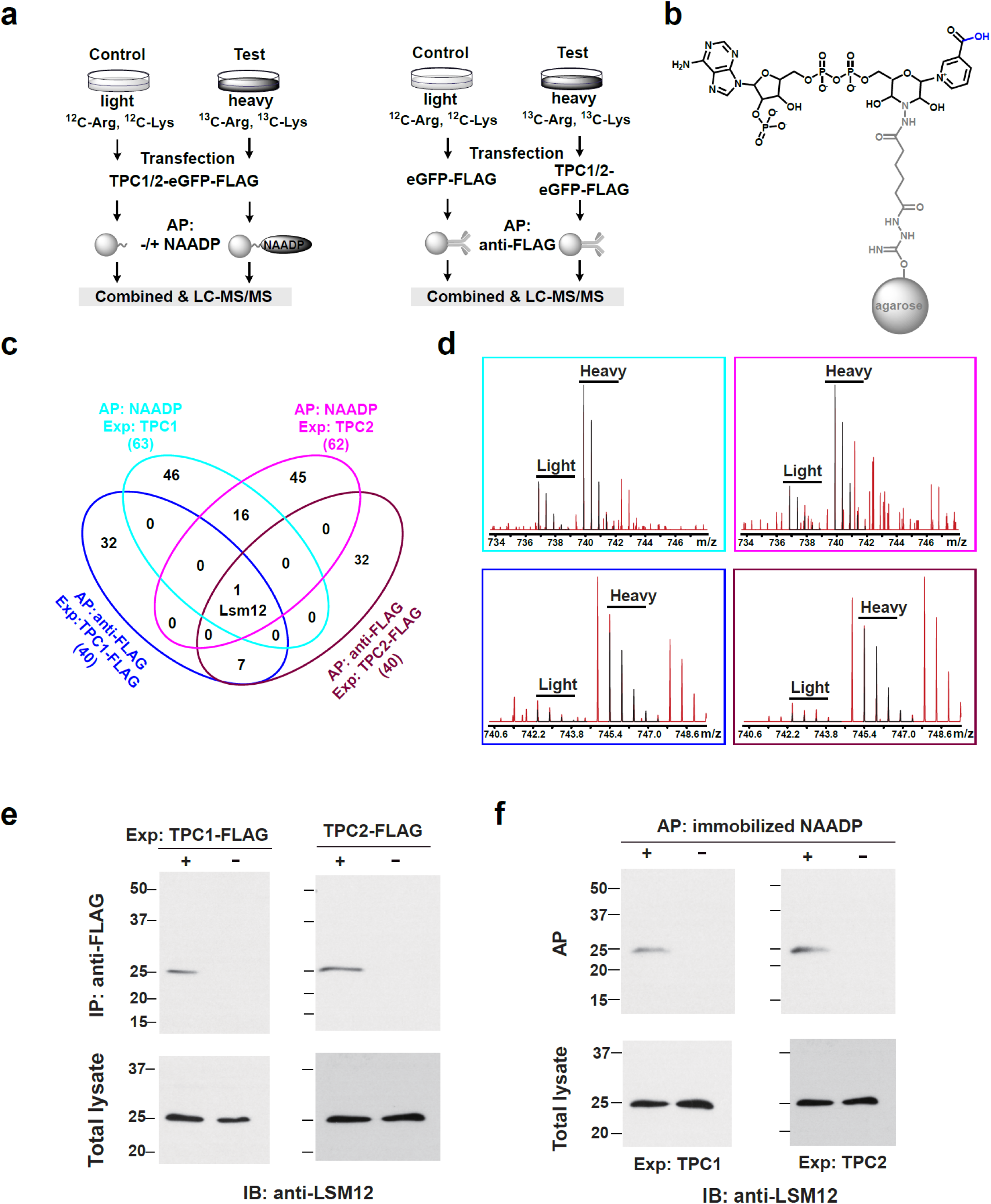
Identification of Lsm12 as a putative NAADP receptor. **a**, Strategies of affinity precipitation- and SILAC-based quantitative proteomic analyses of interacting proteins of NAADP (left) and TPCs (right). **b**, Chemical structures of immobilized NAADP. **c**, Numbers of overlapping and non-overlapping putative interacting proteins identified by MS in the indicated four types of affinity-precipitated samples. **d**, Representative heavy-light peak pairs (black) of Lsm12 peptides. Peptide SQAQQPQKEAALS^194^ (top) was identified in the samples affinity-purified by immobilized NAADP from TPC1- and TPC2-expressing cells. Peptide LQGEVVAFDYQSK^37^ was identified in the TPC1 and TPC2 complexes affinity-purified by an anti-FLAG antibody. **e**, Immunoblot of endogenous Lsm12 in samples immunoprecipitated by an anti-FLAG antibody from cells expressing FLAG-tagged TPC1 or TPC2. **f**, Immunoblot of endogenous Lsm12 in samples pulled down by immobilized NAADP from TPC1- and TPC2-expressing cells. AP, affinity-precipitation. Exp., transient expression. IB, immunoblot.

### Lsm12 is Required and Directly Involved in NAADP-evoked Ca^2+^ Release

To determine the role of Lsm12 in NAADP signaling, we generated a HEK293 Lsm12-knockout (KO) cell line by using the CRISPR/Cas9 genome editing method (**Fig. S3a**). The *Lsm12* alleles in the Lsm12-KO cells have either 1 bp deletion or 68 bps insertion at the start of Lsm12’s exon 3 (**Fig. S3b**). Both mutations result in framing errors after the amino acid residue S46 and thus a truncation of 149 amino acid residues (195 residues in full length). Immunoblotting with an anti-Lsm12 antibody, which recognizes the C-terminal region (**Fig. S3c**), showed that Lsm12 expression was undetectable in the cell lysate of Lsm12-KO cells (**Fig. 2a**). Immunofluorescence staining with the same anti-Lsm12 antibody showed that the staining was broadly present in cytoplasm in WT cells but became invisible in Lsm12-KO cells (**Fig. 2a**). The transiently expressed exogenous recombinant Lsm12 (Myc and FLAG-tagged) replenished Lsm12 expression in Lsm12-KO cells (**Fig. 2a**).

**Figure 2.**
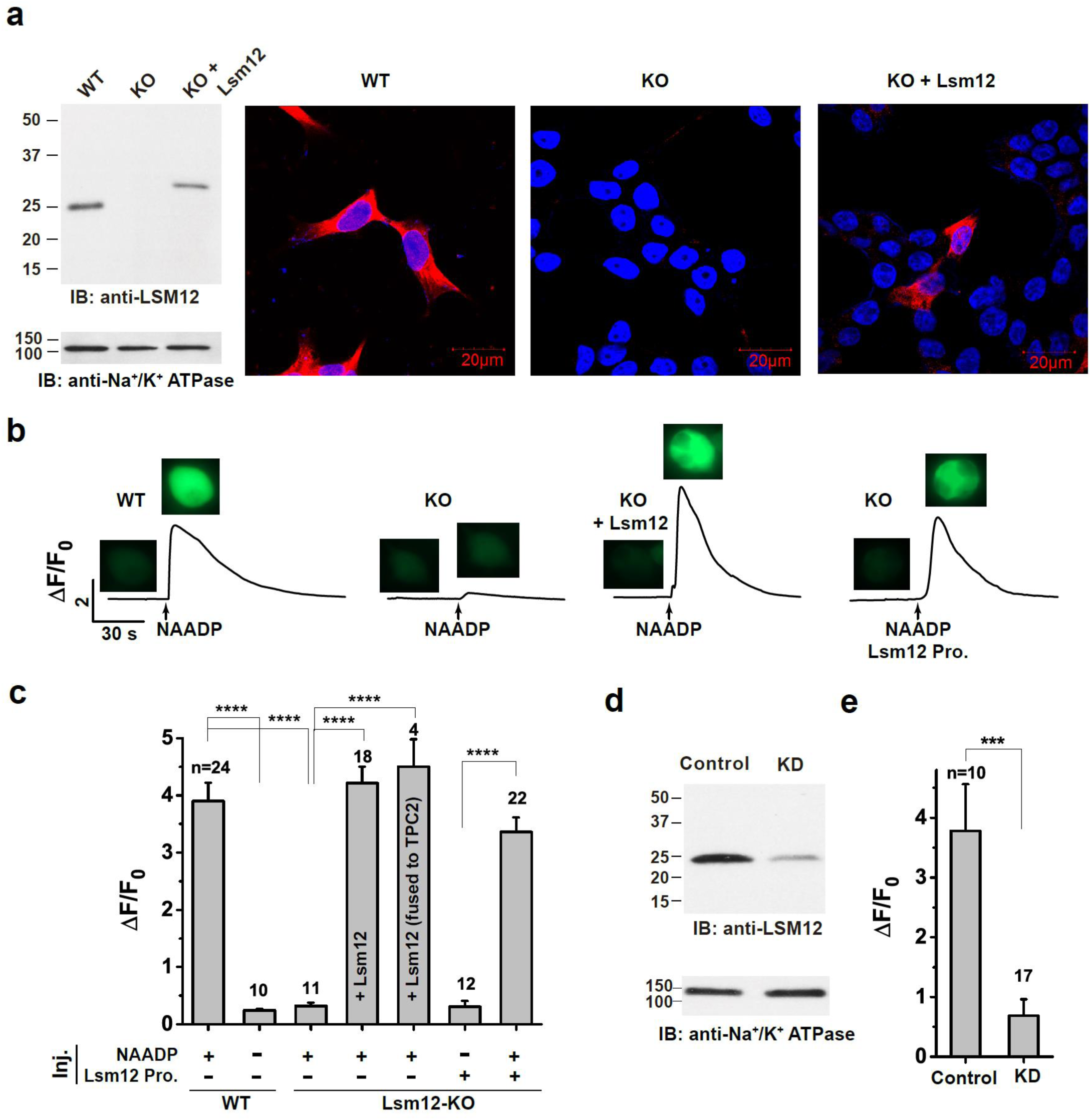
Lsm12 is essential for NAADP-evoked Ca^2+^ release. **a**, Immunoblot and immunofluorescence assays show the loss of Lsm12 expression in the Lsm12-KO cell line. Lsm12 and cell nuclei are shown in red and blue, respectively. **b**, Time course and cell images of NAADP-induced change in fluorescence of Ca^2+^ indicator in TPC2-expressing HEK293 WT and Lsm12-KO cells with/without exogenous Lsm12 expression or protein injection. Cell images were taken before NAADP injection and at the peak of Ca^2+^ increase after injection. **c**, Averaged NAADP-induced changes in Ca^2+^ indicator fluorescence in TPC2-expressing HEK293 WT and Lsm12-KO cells. Exogenous Lsm12 was introduced to cells by either plasmid transfection (labeled in column) or protein injection (labeled at the bottom). **d**, Immunoblot of Lsm12 in cell lysates of SK-BR-3 control and Lsm12-KD cells. **e**, Averaged NAADP-induced changes in Ca^2+^ indicator fluorescence in TPC2-expressing SK-BR-3 control and Lsm12-KD cells. The number of recorded cells (n) for each group was indicated. *** and **** are for *p* values (Student *t* test) ≤0.001 and 0.0001, respectively. IB, immunoblot. KO, knockout. KD, knockdown. Inj., injection.

To determine the functional role of Lsm12, we transiently expressed TPC2 and performed a Ca^2+^-imaging assay in HEK293 WT and Lsm12-KO cells. Using the genetically encoded ultrasensitive fluorescent Ca^2+^ sensor protein GCaMP6f ^24^ to monitor changes of the intracellular Ca^2+^ level, we observed a great (∼ 4-fold) increase in the fluorescence of the Ca^2+^ sensor upon the microinjection of NAADP at 100 nM in pipette solution (**Fig. 2b, c and Fig. S4a**). In agreement with the previously established characteristics of NAADP-evoked Ca^2+^ release ^6-9^, the NAADP-elicited Ca^2+^ signal was largely abolished in the absence of exogenous TPC2 expression or after cells had been pretreated with trans-Ned-19 (10 µM), an antagonist of NAADP-mediated Ca^2+^ release ^25^, or bafilomycin A1 (1 µM), an inhibitor of the endolysosomal H^+^-ATPase that prevents pH-dependent Ca^2+^ accumulation in endolysosomal stores (**Fig. S4b**). Notably, the NAADP-elicited Ca^2+^ increase was fully abolished in TPC2-expressing Lsm12-KO cells as the Ca^2+^ changes were similar to those observed without NAADP (vehicle only) in TPC2-expressing WT cells (**Fig. 2b, c**). The NAADP-evoked Ca^2+^ release was restored in Lsm12-KO cells by co-transfection of TPC2 with Lsm12 (**Fig. 2b, c**) or by expression of Lsm12 as a fused protein tagged to the C-terminus of TPC2 (**Fig. 2c**). In contrast, knockout of Lsm12 expression had no influence on the intracellular Ca^2+^ elevation induced by extracellular ATP (**Fig. S4c**), a purinergic receptor agonist that triggers IP3 production and Ca^2+^ mobilization from the ER. Thus, we conclude that Lsm12 is specifically required for NAADP-evoked Ca^2+^ release.

To determine whether Lsm12 is directly involved in NAADP signaling, we obtained a purified recombinant human Lsm12 protein sample (hLsm12-His_E.coli_) by expressing a 6×His -tagged human Lsm12 protein in *Escherichia coli* and purifying it with immobilized metal affinity chromatography. Notably, microinjection of the purified hLsm12-His_E.coli_ protein (∼ 4 µM in pipette) together with NAADP immediately restored the NAADP-evoked Ca^2+^ release in Lsm12-KO cells (**Fig. 2b, c**). Thus, Lsm12 directly participates in the process of NAADP-evoked Ca^2+^ release in HEK293 cells.

To examine whether Lsm12 is also important for NAADP signaling in other cell lines, we used siRNA to knock down Lsm12 expression in SK-BR-3 cells, a breast cancer cell line that was previously shown to be responsive to NAADP ^7^. We observed that Lsm12 expression at the protein level was greatly reduced in the Lsm12-knockdown (KD) cells (**Fig. 2d**). Correspondingly, the NAADP-evoked Ca^2+^ elevation was also markedly decreased in the Lsm12-KD cells (**Fig. 2e**).

### Lsm12 Directly Binds to NAADP with High Affinity

To determine whether Lsm12 is truly an NAADP receptor, we performed competition ligand binding assays to examine its binding affinity to NAADP. A key feature of the unknown NAADP receptor in NAADP-evoked Ca^2+^ release is its high and specific affinity to NAADP relative to the closely related NADP ^6,26^. To exclude the possibility that some endogenous Lsm12-interacting protein of HEK293 cells might be involved in mediating the observed binding of NAADP to Lsm12, we performed binding assays using the purified hLsm12-His_E.coli_ protein that was produced in a prokaryotic expression system and confirmed to be functional in NAADP signaling upon microinjection of the protein into HEK293 Lsm12-KO cells (**Fig. 2b, c**). We observed that hLsm12-His_E.coli_ can be effectively pulled down by immobilized NAADP and that the formation of immobilized NAADP−Lsm12 complex was competitively prevented by inclusion of low concentrations (nM range) of free NAADP (**Fig. 3a and Fig. S5a**). The estimated K_d_ for NAADP binding is ∼30 nM, whereas NADP up to 100 µM had nearly no effect on the binding between hLsm12-His_E.coli_ and immobilized NAADP (**Fig. 3a and Fig. S5a**). Since ^32^P-NAADP-based radioligand binding was the common method used in determining the binding affinity of NAADP to endogenous receptors in HEK293 cells ^6,26,27^, we synthesized ^32^P-NAADP from ^32^P-NAD as previously described ^27^ and purified ^32^P-NAADP with thin-layer chromatography (**Fig. S5b**). We observed that ^32^P-NAADP can effectively bind to hLsm12-His_E.coli_ and that the binding can be competitively blocked by preincubation of the protein with nM concentrations of non-radiolabeled NAADP (**Fig. 3b**). The ^32^P-NAADP binding curve can be fitted with a high affinity site (*K*_d_ = ∼20 nM; 79% in fraction) and a low affinity site (*K*_d_ = ∼5 µM; 21% in fraction) (**Fig. 3b**). NADP up to 100 μM showed nearly no influence on the binding of ^32^P-NAADP to hLsm12-His_E.coli_ (**Fig. 3b**). These competition ligand binding assays demonstrate that Lsm12 is a direct high-affinity receptor for NAADP.

**Figure 3.**
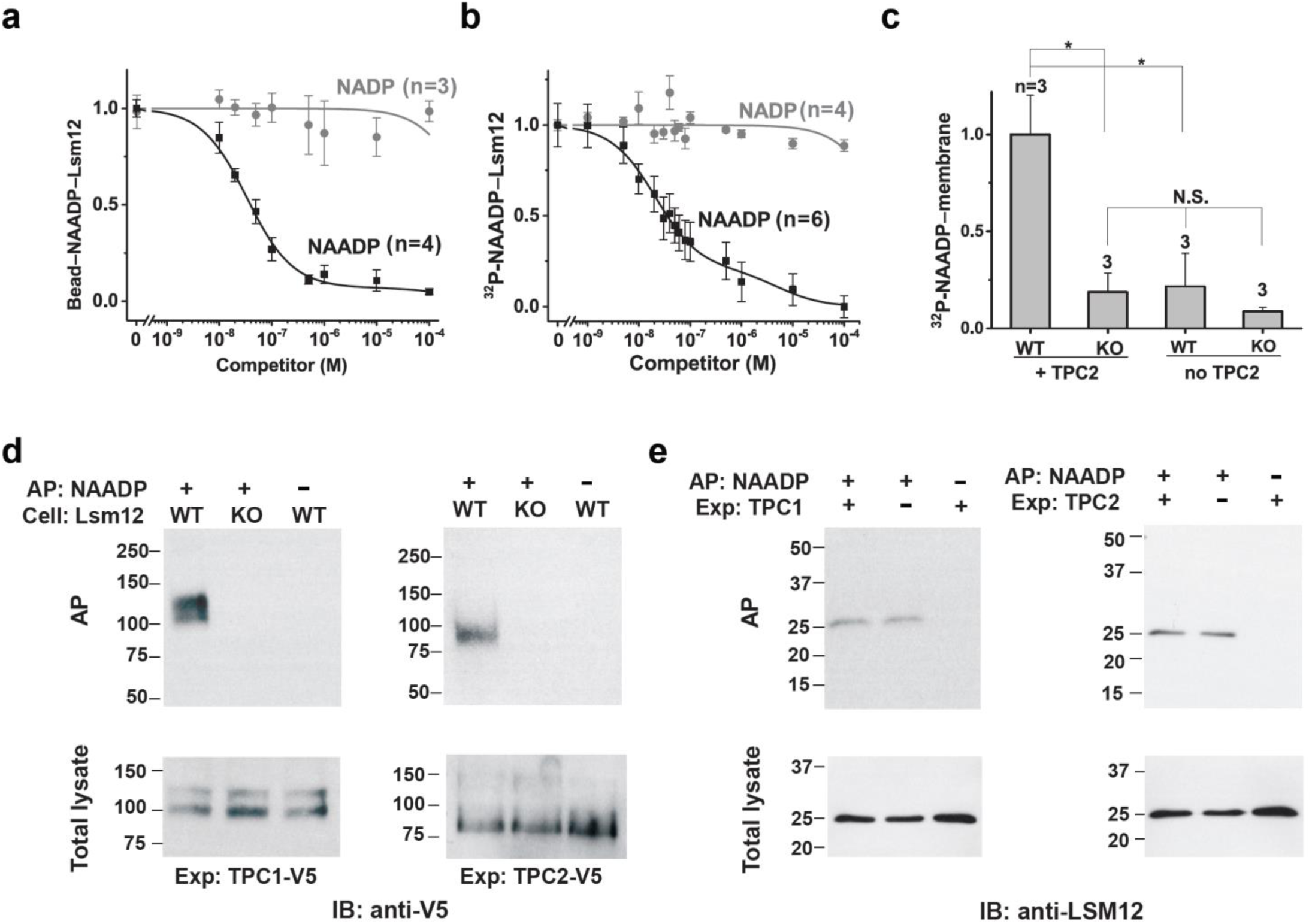
Lsm12 functions as a high affinity NAADP receptor. **a**, Plot of the relationship between hLsm12-His_E.coli_ pulled down by immobilized NAADP and the concentrations of free NAADP and NADP. **b**, Competition radioligand binding assay of the association between ^32^P-NAADP and purified hLsm12-His_E.coli_ in the absence or presence of various concentrations of non-radiolabeled NAADP and NADP. **c**, Specific binding of ^32^P-NAADP to cell membranes in WT and Lsm12-KO cells with/without exogenous TPC2 expression. **d**, Immunoblot of TPC1 and TPC2 pulled down by immobilized NAADP from TPC1- and TPC2-expressing HEK293 WT and Lsm12-KO cells. **e**, Immunoblot of Lsm12 pulled down by immobilized NAADP from cells with/without exogenous TPC1 and TPC2 expression. AP, affinity precipitation. Exp, expression. AP: NAADP +/–, AP with/without immobilized NAADP. * is for *p* values (Student *t* test) ≤0.05.

### Lsm12 Mediates the Apparent Association of NAADP to TPCs and TPC-containing Membranes

The high binding affinity between NAADP and hLsm12-His_E.coli_ is comparable to the previously reported binding affinity between NAADP and TPC-enriching protein or membrane preparations ^6,26^. We reasoned that the apparent association between NAADP and TPCs or TPC-containing cell membranes might result from Lsm12, which brings NAADP and TPCs together. To exam this possibility, we first determined the contribution of Lsm12 in NAADP binding to TPC2-expressing cell membranes using a competition radioligand binding assay as previously described ^6,26^. We preincubated cell membranes of TPC2-expressing cells with different concentrations of non-radiolabeled NAADP before the addition of ^32^P-NAADP (∼1 nM). The specific binding to the cell membranes was estimated by subtracting the bound total ^32^P from that of non-specifically bound ^32^P determined in the presence of 100 μM non-radiolabeled NAADP. Similar to the previous report ^6^, we observed a high affinity ^32^P-NAADP binding of TPC2-expressing membranes that was competed off by 40−75% with the preincubation of 5−50 nM non-radiolabeled NAADP (**Fig. S5c**). However, in membranes prepared from TPC2-expressing Lsm12-KO cells, the specific binding of ^32^P-NAADP was mostly lost (e.g., by ∼80% in the absence of non-radiolabeled NAADP) (**Fig. 3c and Fig. S5c**). Second, given that Lsm12 is a cytosolic protein (**Fig. 2a**), an Lsm12 interacting protein on the membrane is expected to be needed for the apparent association of NAADP to cell membranes. We found that TPC2 expression was also required for the association of ^32^P-NAADP to cell membranes, suggesting that Lsm12 is mostly associated with the membrane through its interaction with TPCs (**Fig. 3c**). Third, both TPC1 and TPC2 were pulled down by immobilized NAADP (**Fig. 3d**) but only in the presence of Lsm12 as immobilized NAADP pulled down neither TPC2 nor TPC1 from the Lsm12-KO cells (**Fig. 3d**). In contrast, immobilized NAADP pulled down Lsm12 in a largely TPC expression–independent manner since no significant difference was observed in the absence or presence of exogenous TPC1 or TPC2 expression in HEK293 cells (**Fig. 3e**). These results demonstrate that TPCs are not NAADP receptors and that the apparent association of NAADP to TPCs or TPC-containing membranes was actually mediated by Lsm12.

### Lsm12 Mediates NAADP Signaling via Its Lsm Domain

Lsm12 possesses an N-terminal Lsm domain and a putative C-terminal anticodon-binding (AD) domain ^28^ (**Fig. S6a**). To determine which domain or region of Lsm12 is involved in NAADP-evoked Ca^2+^ release, we generated three truncation mutants ΔLsm, ΔAD, and Δlinker by deletion of the Lsm domain, the C-terminus from the putative AD domain to the C-terminal end, and the linker region between the Lsm and AD domains, respectively. Upon transfection of the constructs into the Lsm12-KO cells, we observed that the ΔLsm mutant was unable to rescue the NAADP-evoked Ca^2+^ release, whereas the ΔAD and Δlinker mutants remained largely functional in NAADP signaling (**Fig. 4a**). To understand the function of the Lsm domain, we tested the mutants’ function in binding to NAADP and TPC2. We found that the ΔLsm mutation eliminated the capability of the protein’s binding to immobilized NAADP, whereas ΔAD and Δlinker mutations had no such effect (**Fig. 4b**). Similarly, the ΔLsm mutation abolished the association between ^32^P-NAADP and a recombinant FLAG-tagged Lsm12 protein that was heterologously expressed in Lsm12-KO cells and immunoprecipitated by an anti-FLAG antibody (**Fig. 4c**). By co-immunoprecipitation, we also found that only ΔLsm caused a loss of the association between Lsm12 and TPC2 (**Fig. 4d**). Thus, the Lsm12’s Lsm domain plays a dual role by interacting with both NAADP and TPC2. However, NAADP and TPC2 must interact with different regions of the Lsm domain since the pulldown of Lsm12 by immobilized NAADP was similar in the absence and presence of TPC1 and TPC2. (**Fig. 3e**). Furthermore, no significant difference in the co-immunoprecipitation of Lsm12 and TPC2 was observed in the absence and presence of NAADP (**Fig. 4e**).

**Figure 4.**
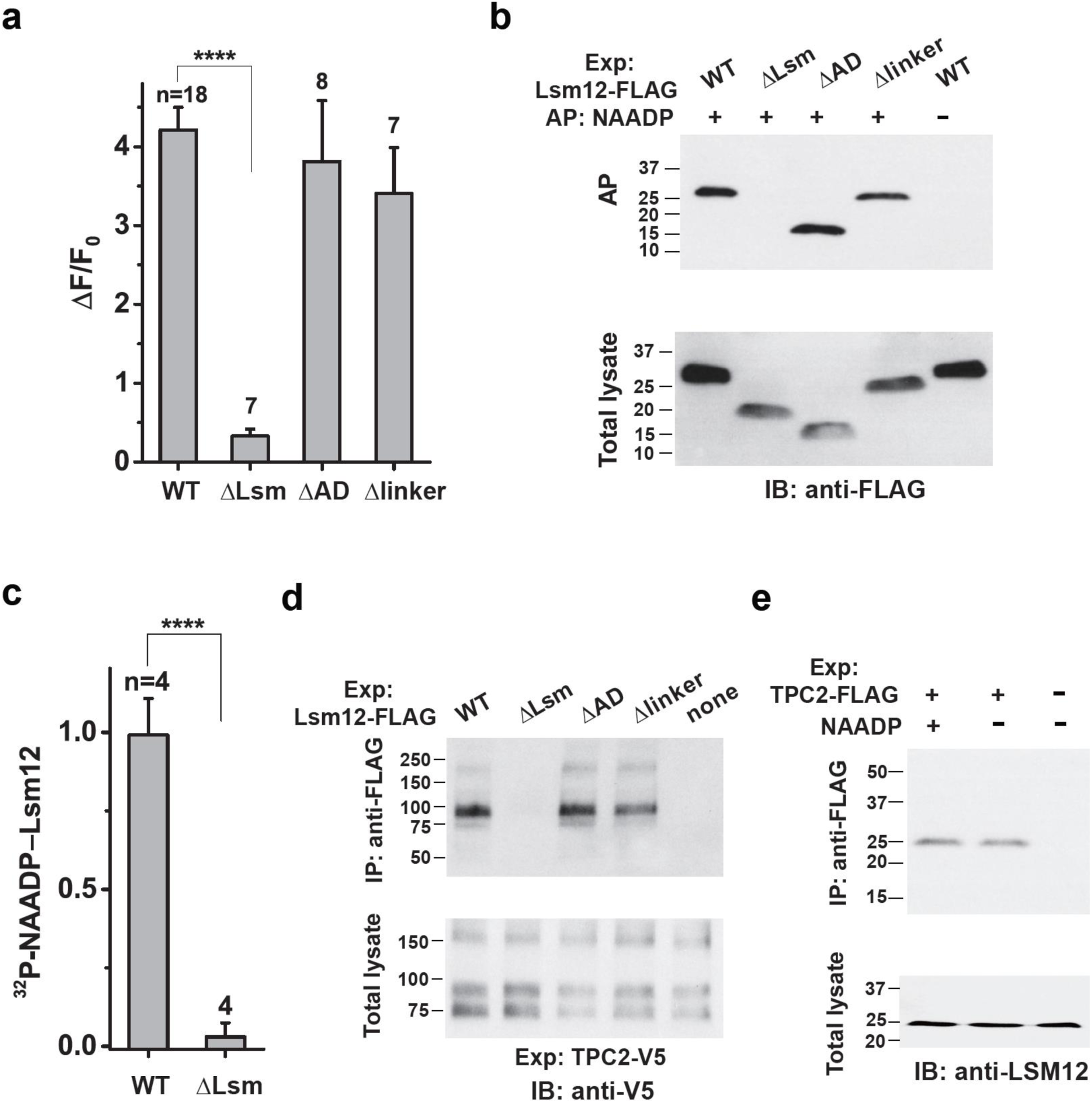
Lsm12 functions in NAADP signaling via its Lsm domain. **a**, Averaged NAADP-induced changes in Ca^2+^ indicator fluorescence in Lsm12-KO cells transfection with Lsm12 WT and mutant constructs. The number of recorded cells (n) for each group was indicated. **b**, Immunoblot of Lsm12 WT and mutants pulled down by immobilized NAADP. **c**, Specific binding between ^32^P-NAADP and FLAG-tagged Lsm12 WT and ΔLsm proteins that were transiently expressed in Lsm12-KO cells and immunoprecipitated by an anti-FLAG antibody. **d**, Co-immunoprecipitation between TPC2 and Lsm12 mutants. **e**, Co-immunoprecipitation between TPC2 and Lsm12 in the absence or presence of 100 µM NAADP. Exp, expression.

### Lsm12 Is Required for NAADP-induced TPC2 activation

TPCs have been considered as the key lysosomal cation channels responsive to NAADP stimulation for intracellular Ca^2+^ release. However, controversy exists as the TPCs’ sensitivity to NAADP was not observed in multiple reports ^16-19^. Similarly, we were also unable to detect any NAADP-activated TPC2 currents under more invasive recording conditions by either a “caesarean section”-like acute isolation of enlarged lysosomes in a whole lysosome recording of WT channels or by excision of plasma membrane in an inside-out recording of the plasma membrane-targeted mutant (TPC2^L11A/L12A^) ^8^ channels. To retain the NAADP sensitivity of the channels, we measured NAADP microinjection-induced whole-cell currents of voltage-clamped HEK293 cells that expressed TPC2^L11A/L12A^ (**Fig. 5a**). With this novel and less invasive method that matches our experimental condition of NAADP microinjection-evoked Ca^2+^ release, we observed robust NAADP-evoked Na^+^ currents (inward) which were ∼ 0.5 nA or 38.7 ± 4.8 pA/pF on average at -120 mV for an injection solution containing 1 μM NAADP (**Fig. 5a, b**). The recorded currents were slightly smaller (∼ 0.3 nA or 24.6 ± 5.4 pA/pF at -120 mV) when the injection solution contained 100 nM NAADP (**Fig. 5b**), a concentration that can achieve maximal NAADP-evoked Ca^2+^ release in our Ca^2+^-imaging experiments (**Fig. S4a**). Such currents were mostly absent when no exogenous TPC2 was expressed (**Fig. 5a, b**), when the lysosome-targeted WT TPC2 was used (**Fig. 5b**), or in the presence of a dominant-negative mutation L265P ^8^ (**Fig. 5b**), indicating that the NAADP-evoked inward Na^+^ currents were mostly originated from the plasma membrane-targeted TPC2^L11A/L12A^ channels. Consistent with Lsm12 as the NAADP receptor, the NAADP-induced TPC currents in TPC2-expressing Lsm12-KO cells were absent. The response was restored by supplementing Lsm12 via transfection of a Lsm12-expressing plasmid, fusion of the Lsm12 to the C-terminus of TPC2, or co-microinjection of NAADP with the purified Lsm12 protein (**Fig. 5a, b**). Microinjection of the purified Lsm12 protein alone without NAADP didn’t result in a significant increase in the inward Na^+^ currents (**Fig. 5b**), suggesting that both NAADP and Lsm12 are required for TPC2 activation. As it is expected for the function of Lsm12 as an NAADP receptor and an interacting protein of TPCs, these results demonstrate that Lsm12 is required to mediate TPC2 activation by NAADP.

**Figure 5.**
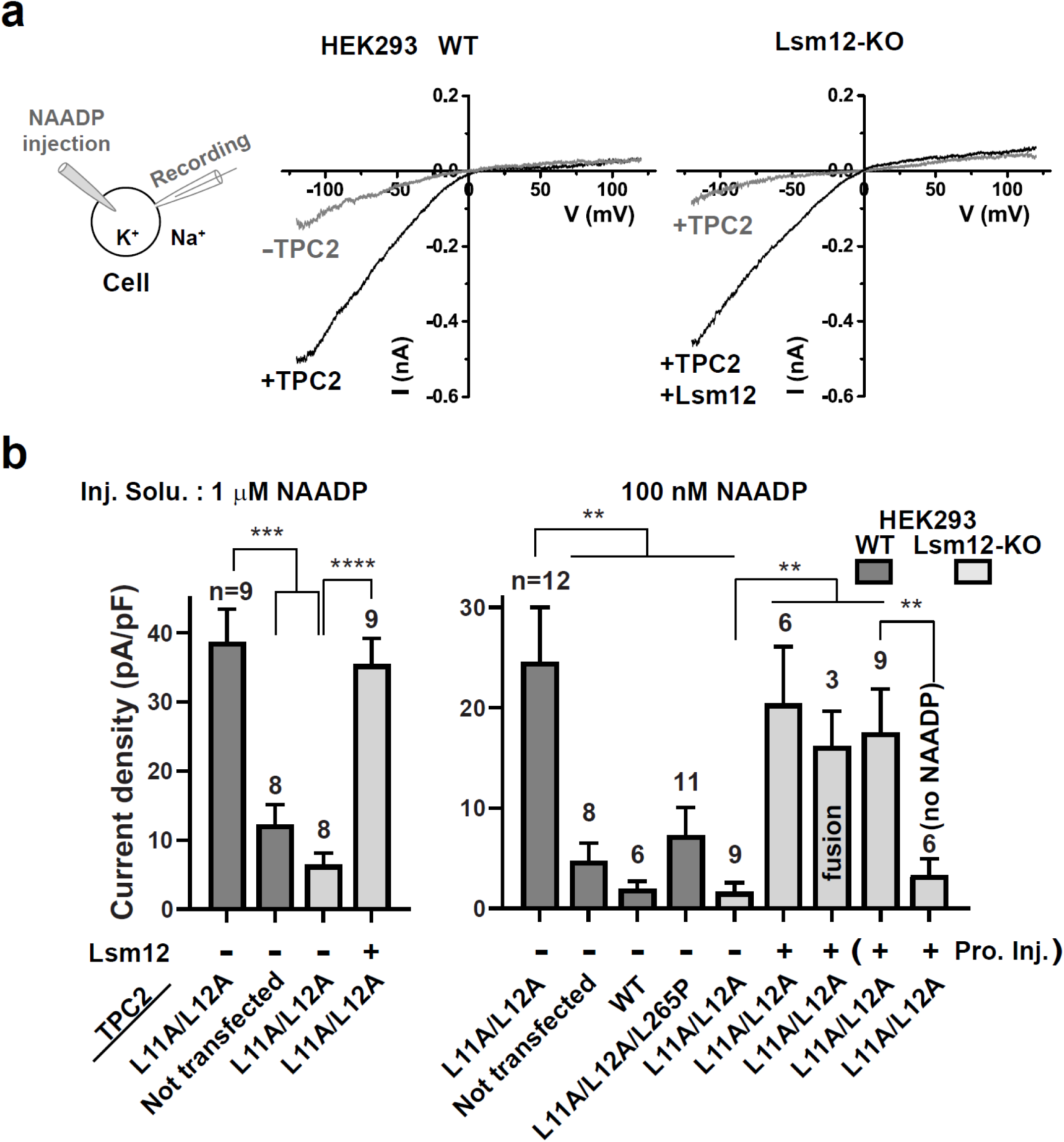
Lsm12 is required for TPC2 activation by NAADP. **a**, Averaged traces of NAADP (1 μM in pipette solution) microinjection-induced whole cell currents (averaged) in HEK293 WT and Lsm12-KO cells transfected with/without TPC2^L11A/L12A^ and Lsm12. **b**, Averaged current density of NAADP microinjection-induced whole cell currents in HEK293 WT and Lsm12-KO cells. The cells were transfected with/without Lsm12 and TPC2 (WT or mutants). NAADP was injected at 1 μM (left) or 100 nM (right) and the purified Lsm12 protein (last 2 columns on the right) was injected at ∼ 100 ng/ml (∼ 4 μM) in injection pipette solutions. The number of recorded cells (n) for each group was indicated. **, *** and **** are for *p* values (Student *t* test) ≤0.01, 0.001, and 0.0001, respectively. Inj. Solu., injection solution. Pro. Inj., protein injection.

### Lsm12 Is Important for NAADP-signaling in Mouse Embryonic Fibroblasts

To evaluate the importance of Lsm12 in primary cells, we generated Lsm12 mutant mice with the CRISPR/Cas9 method. The ablation of Lsm12 function by a frameshifting mutation, a deletion of 18 bps (ATGTCCCTCTTCCAGTGG) in the 3^rd^ exon, appeared to be embryonically lethal as no homozygous mouse could be obtained. Thus, a mutant mouse harboring a different 18 bp (TCTTCCAGTGGAAACCC) deletion in the 3^rd^ exon was used. This non-frameshifting mutation, designated as Lsm12^Δ45-50^, results in a deletion of 6 residues (SSSGKP^50^) in the Lsm domain. Given that mouse embryonic fibroblasts (MEFs) were frequently used in studying NAADP signaling or TPC function in primary cells ^11,29-31^, we isolated MEFs from WT and Lsm12^Δ45-50^ mice and measured the cells’ response to NAADP. We observed that the NAADP-evoked Ca^2+^ elevation was greatly reduced by 75% in MEFs of the Lsm12^Δ45-50^ mice as compared to that of WT mice (**Fig. 6a**). When the Lsm12^Δ45-50^ mutant construct was transiently expressed in the HEK293 Lsm12-KO cells, it was also much less effective (a reduction by ∼ 60%) than WT Lsm12 in rescuing the NAADP-evoked Ca^2+^ release in the HEK293 Lsm12-KO cells (**Fig. 6b**). Consistently, the NAADP-induced TPC2 currents were not effectively rescued by the Lsm12^Δ45-50^ mutant (∼ 60% reduction in efficiency as compared to WT) upon its transient expression in HEK293 Lsm12-KO cells (**Fig. 6c**). The deficiency of the Lsm12^Δ45-50^ mutant in supporting NAADP-induced Ca^2+^ release and TPC2 current activation was not caused by reduced expression as the mutant and WT were similarly expressed in MEFs (**Fig. 6d**) and the mutant displayed a higher transient expression level than WT in HEK293 Lsm12-KO cells (**Fig. 6e**). The latter could result in an underestimate of the mutant’s deficiency in HEK293 cells. These results obtained with MEFs and HEK293 cells show deficiency of the Lsm12^Δ45-50^ mutant in NAADP signaling and support a key role of Lsm12 in NAADP signaling in MEFs.

**Figure 6.**
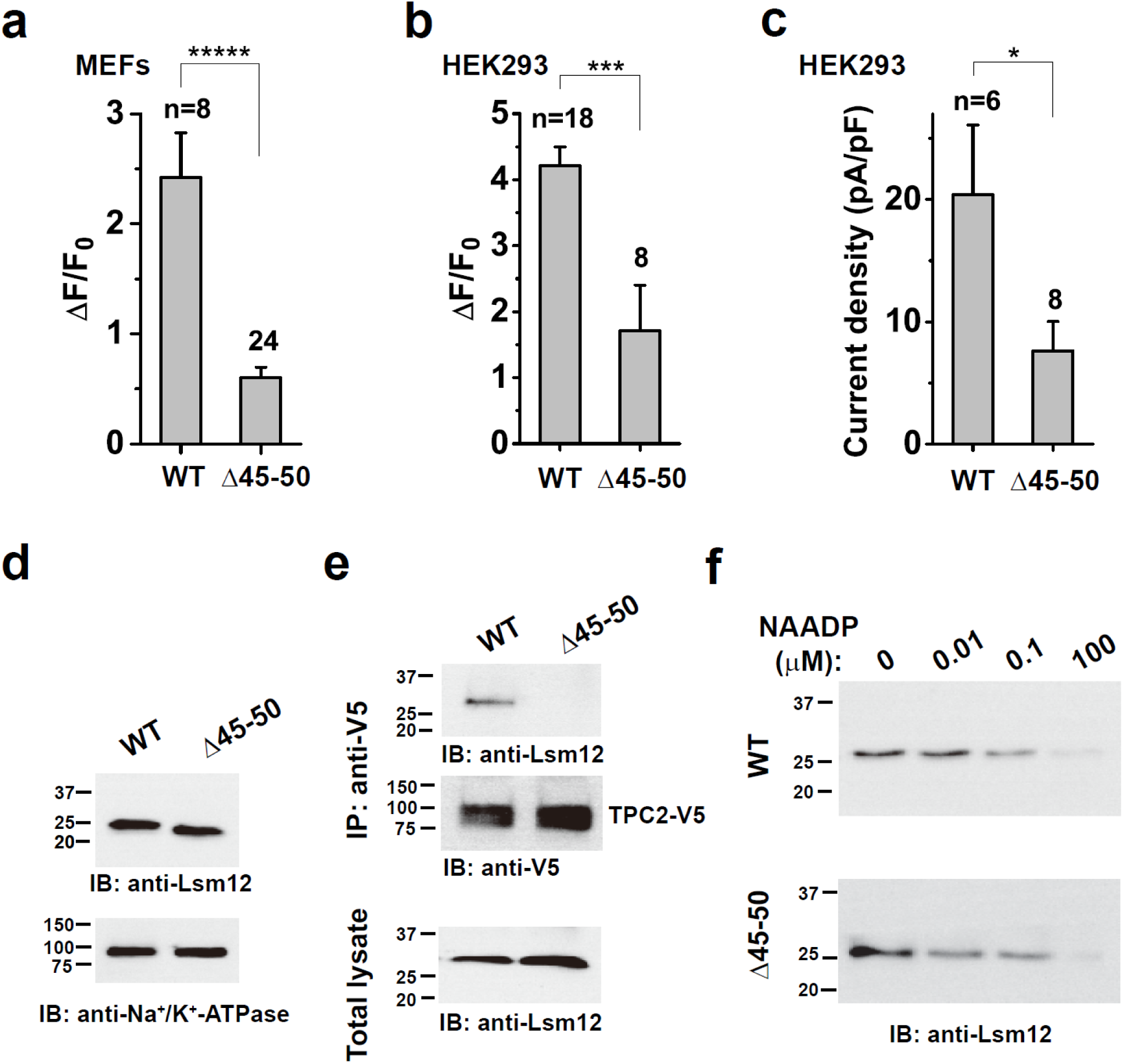
Lsm12 function in NAADP signaling is compromised by Δ45-50 mutation in MEFs and HEK293 cells. **a**, Averaged NAADP-induced changes in Ca^2+^ indicator fluorescence in MEFs prepared from WT and Lsm12^Δ45-50^ mutant mice. **b**, Averaged NAADP-induced changes in Ca^2+^ indicator fluorescence in HEK293 Lsm12-KO cells transiently expressing TPC2 and Lsm12 WT or Δ45-50 mutant. **c**, Averaged current density of NAADP (100 nM in pipette solution) microinjection-induced whole cell currents in HEK293 Lsm12-KO cells transiently expressing TPC2 and Lsm12 WT or Δ45-50 mutant. **d**, Immunoblot of Lsm12 in MEFs prepared from WT and Lsm12^Δ45-50^ mutant mice. MEFs were prepared and pooled from 5 WT and 6 mice Lsm12^Δ45-50^ mutant mice respectively. **e**, Co-immunoprecipitation (IP) of TPC2 (V5-tagged) and LSM12 WT or Δ45-50 mutant transiently expressed in HEK293 LSM12-KO cells. **f**, Immunoblot of Lsm12 WT and Δ45-50 mutant that were transiently expressed in HEK293 Lsm12-KO cells and pulled down by immobilized NAADP in the presence of different concentrations of free NAADP. The number of recorded cells (n) for each group was indicated. **\***, ***, and ***** are for *p* values (Student *t* test) ≤0.05, 0.001 and 0.00001, respectively. IB, immunoblot. IP: immunoprecipitation.

To understand the mechanisms underlying the Lsm12^Δ45-50^ mutant’s defects in NAADP-induced Ca^2+^ release and TPC2 activation, we assayed its functions in TPC and NAADP bindings. We found that its association with TPC2 is compromised. Co-immunoprecipitation of the co-expressed TPC2 and Lsm12 showed that the Δ45-50 mutant was barely pulled-down together with TPC2 (**Fig. 6e**). The Lsm12^Δ45-50^ mutant remained functional in its binding with NAADP as 10 nM and 100 nM NAADP still competitively reduced the protein’s association with immobilized NAADP in a manner similar to that observed with the WT protein (**Fig. 6f**). These suggest that the complex formation between Lsm12 and TPC2 is likely required in NAADP-induced TPC2 activation and Ca^2+^ mobilization from intracellular stores.

## DISCUSSION

The unresolved molecular identity of the NAADP receptor has limited our understanding of the molecular basis and mechanisms of NAADP-evoked Ca^2+^ release and signaling. The present study has identified a previously largely uncharacterized protein, Lsm12, as an NAADP receptor essential for NAADP-evoked Ca^2+^ release. Lsm12 has a predicted protein size of 21.7 kDa and an observed protein size of ∼25 kDa, which is close to the previously reported doublet protein band sizes (22/23 kDa) of putative NAADP receptors photolabeled by [^32^P]5-azido-NAADP in cell lysates^20^. According to The Human Protein Atlas (www.proteinatlas.org), Lsm12 at the RNA level is ubiquitous and abundantly expressed in different tissues and cell lines. Given that Lsm12 is functionally critical for NAADP-evoked Ca^2+^ release in all 3 tested cell types (HEK293, SK-BR-3 and MEFs), we anticipate that Lsm12 as an NAADP receptor plays a universal role in NAADP signaling in different cells.

Lsm12 belongs to the Lsm protein family, which includes smaller Lsm proteins (Lsm1-10) that contain only an Lsm domain and the larger Lsm proteins (Lsm11, 12, 14A, 14B, and 16 in human) that contain an extra C-terminal non-Lsm domain ^28^. Although much less or little is known about the functions of larger Lsm proteins, smaller Lsm proteins (e.g., Lsm1–8) generally function as scaffolds or chaperones to bind to RNA oligonucleotides, facilitating RNA assembly, modification, storage, transportation, or degradation and thus globally affecting cell gene expression ^32^. In spite of its potential RNA processing function, we have observed no evidence that Lsm12 can indirectly affect NAADP signaling by transcriptional regulation of other proteins that might be relevant in NAADP signaling (e.g., TPCs for Ca^2+^ release from lysosomes and ryanodine receptors for Ca^2+^-release from ER). We have found that exogenous TPC1 and TPC2 were similarly expressed in HEK293 WT and Lsm12-KO cells (**Fig. 3d**). Furthermore, the extracellular ATP-induced Ca^2+^ mobilization from the ER was not affected by Lsm12 (**Fig. S4c**).

The Lsm domain is predicted to form a highly conserved tertiary structure of Lsm fold that consists of a 5-stranded anti-parallel β-sheet (**Fig. S6b**). As seen in many other Lsm proteins, the Lsm domain may oligomerize into a Lsm ring that allows binding to oligonucleotide (adenine or uracil) ^32^. Given that NAADP is essentially a type of dinucleotide, Lsm12 is well positioned to serve as an NAADP receptor via its Lsm domain. Although many other Lsm proteins are also well expressed at the RNA level in different cells (www.proteinatlas.org), Lsm12 is likely a dominant NAADP receptor as no other Lsm protein has been identified in our proteomic analysis of NAADP binding proteins in HEK293 cells (**Table S1**) and Lsm12 was required in NAADP-evoked Ca^2+^ release in different cells.

Since the reports of a key role of TPCs in NAADP signaling ^6,7^, it had been controversial whether TPCs are NAADP receptors ^20,22^ and whether TPCs can be activated by NAADP ^16-19,22^. Our identification of Lsm12 as an NAADP receptor and characterization of its relationship to TPCs have clearly consolidates the notion that TPCs are not direct receptors of NAADP ^20-22^. In addition, we demonstrated NAADP-induced TPC2 currents under a less invasive patch clamp recording condition with a combination of whole cell recording and NAADP-microinjection. Furthermore, the physical association between Lsm12 and TPC2 likely plays a role in NAADP signaling as the Lsm12^Δ45-50^ mutant, which has a weakened association with TPC2, was deficient in NAADP-evoked TPC2 activation and intracellular Ca^2+^ elevation. These led us to conclude that TPCs are regulated by NAADP via the receptor Lsm12 (**Fig. S6c**). Further studies will be needed to elucidate the mechanisms underlying the TPC activation caused by the NAADP binding to Lsm12. Overall, our studies reveal the molecular identity of the NAADP receptor to be a Lsm protein (Lsm12) and establish its essential role in NAADP-evoked TPC2 activation and Ca^2+^ mobilization from intracellular stores. Our findings thus provide a new molecular basis toward elucidating the mechanisms and function of NAADP signaling.

## METHODS

### Mammalian cells, culture, and manipulation of protein expression

A variant of the HEK293 cell line, 293H (Invitrogen), which has better adherence in monolayer culture, was used throughout the study. SK-BR-3 cells were obtained from ATCC. HEK293 and SK-BR-3 cells were cultured in Dulbecco’s modified Eagle’s medium and RPMI 1640 medium, respectively, supplemented with 10% fetal bovine serum, 50 units/ml penicillin, and 50 mg/ml streptomycin. A Lsm12-KO cell line of HEK293 cells was generated with Synthego’s chemically modified sgRNA 5’-CCAGAAUGUCCCUCUUCCAG-3’ and GeneArt Platinium Cas9 nuclease (ThermoFisher Scientific) that were transfected together into cells using Lipofectamine CRISPRMAX Cas9 transfection reagent (ThermoFisher Scientific).

Knockdown of Lsm12 expression in SK-BR-3 was achieved with a predesigned dicer-substrate siRNA (DsiRNA) (Cat# hs.Ei.LSM12.13.2 from IDT) targeting exon 3 of Lsm12. A non-targeting DsiRNA (Cat# 51-01-14-03 from IDT) was used as a negative control. Recombinant cDNA constructs of human TPC1, TPC2 with/without eGFP, FLAG, and/or V5 tags were constructed with pcDNA6 vector (Invitrogen). Recombinant human Lsm12-expression plasmids were obtained either commercially (pCMV-Lsm12-Myc-FLAG from OriGene) or were constructed with pcDNA6 (pCDNA6-Lsm12-FLAG-His). Cells were transiently transfected with plasmids with use of transfection reagent of Lipofectamine 2000 (Invitrogen), PolyFect (QIAGEN), or FuGENE HD (Promega) and subjected to experiments within 16-48 h after transfection.

### Quantitative proteomic analyses of NAADP- or TPC-interacting proteins

HEK293 cells were cultured in SILAC-compatible DMEM supplemented with 10% dialyzed fetal bovine serum and 50 mM ^13^C or ^12^C-labeled lysine and arginine (all reagents were from Thermo Fischer Scientific) for at least 6 days to achieve an incorporation efficiency of >90%. Immobilized NAADP and anti-FLAG antibody were used for affinity-precipitation of interacting proteins of NAADP and FLAG-tagged TPCs, respectively.

Immobilized NAADP was prepared as previously reported ^23^. Briefly, 1 μmol of NAADP was incubated with 100 μl of 100 mM sodium periodate/100 mM sodium acetate (pH5.0) mixture in the dark for 1 h. The unreacted sodium periodate was removed by adding 200 mM potassium chloride to the solution, incubating on ice for 5 min, and then centrifuging at 170,000 *g* for 10 min. The supernatant was then incubated with sodium acetate, which was pre-balanced with adipic acid dihydrazide-Agarose beads (Cat# A0802 from Sigma-Aldrich) at 4 °C in the dark for 3 h. The NAADP-conjugated beads were then washed sequentially with 1 M NaCl and phosphate buffered saline (PBS). Anti-FLAG antibody was immobilized to protein-A agarose beads without chemical crosslinking.

For affinity-precipitation, proteins were solubilized from the heavy- and light-labeled cells with a lysis buffer containing 2% dodecyl maltoside in 150 mM NaCl and 20 mM Tris-HCl (pH 7.4). The solubilized proteins were collected as supernatants after centrifugation at 17,000 *g* for 10 min and then incubated overnight with beads of immobilized NAADP or anti-FLAG antibody at 4 °C. After being washed with lysis buffer for 3 times, the bound proteins were eluted with either 100 μg/ml FLAG peptide for collecting interacting proteins of FLAG-tagged TPCs, or 4% SDS for collecting NAADP-interacting proteins.

For SILAC quantification by mass spectrometry, the heavy- and light-labeled eluates were merged. After a brief separation by a short run of SDS-PAGE, the protein-containing gels were excised into three different parts and then subjected to in-gel reduction with dithiothreitol, alkylation with iodoacetamide, digestion with trypsin, and mass spectrometric analysis with a HPLC-tandem mass spectrometer Orbitrap Fusion Lumos Tribrid (ThermoFisher Scientific), as we recently described^33^. Peptides were eluted with a 5% to 50% linear gradient of 0.1% formic acid in 100% acetonitrile for 60 min at a flow rate of 300 nl/min. A full scan was acquired from the range of 300-1800 m/z. The 10 most intense peaks were sequentially isolated for MS/MS experiments with the collision induced dissociation (CID) mode. The spectra labeling, database search, and quantification were carried out with Mascot Distiller (v 2.6) and Mascot (v 2.4) (MatrixScience). A quantification value from a spectra pair is considered valid only when two parameters were met in Mascot Distiller: correlation coefficient between the observed precursor isotope distributions and the predicted ones should be larger than 0.6 and the standard error after the fit to the signal intensities of each precursor pair should be less than 1. Only proteins with heavy:light ratio of ≥3 and a mascot score ≥40 were reported as differential or putative interacting proteins.

### Immunoprecipitation, immunoblot, and immunofluorescence analyses

Immunoprecipitation, immunoblot, and immunofluorescence analyses of protein interactions and expression were performed similarly to analyses we recently reported^33^. The following antibodies were used: anti-FLAG antibodies (Cat# F7425 and F3165 from Sigma-Aldrich), anti-V5 antibody (Cat# R96125 from Invitrogen), and anti-LSM12 antibody (Cat# EPR12282 from Abcam). For immunoprecipitation with anti-FLAG or anti-V5 antibody, FLAG- or V5-peptide was used to elute the antibody-trapped proteins from the beads.

### Expression and purification of recombinant human Lsm12 with *E. coli*

The expression plasmid of recombinant human Lsm12 protein hLsm12-His_E.coli_ was constructed with pET vector and transformed into *E. coli* strain DE3 (BL21). The cells were grown at 37 °C and collected 4 h after induction with 1 mM IPTG. Cells were harvested by centrifugation and broken by sonication or B-PER (Thermo Fisher) in a TBS buffer (150 mM NaCl and 20 mM Tris-HCl pH 8.0). After centrifugation at 12,000 *g* for 10 min, the soluble fraction was collected as supernatant. The 6×His -tagged LSM12 was purified by passing the soluble fraction through a Ni-NTA column and then eluting the column with 300 mM imidazole in TBS. Imidazole in elute was removed by ultrafiltration.

### NAADP binding assays

The NAADP binding assay with immobilized NAADP was performed in a similar manner as that described above for affinity-precipitation of NAADP-interacting proteins except that the NAADP-conjugated beads were preincubated with various concentrations of label-free/unmodified NAADP or NADP before incubation with purified hLsm12-His_E.coli_ (5 nM). The ^32^P-NAADP was synthesized from ^32^P-NAD (Cat# BLU023 from PerkinElmer Co.) following two sequential enzymatic reactions ^27. 32^P-NAD was first catalyzed into ^32^P-NADP with 0.5 U/ml NAD kinase (Cat# AG-40T-0091from Adipogen Co.) in the presence of 5 mM Mg^2+^-ATP and 100 mM HEPES, pH 7.4 at 37 °C for 4 h. The synthesized ^32^P-NADP was converted into ^32^P-NAADP with 1 μg/ml ADP-ribosyl cyclase (Cat# A9106 from Sigma-Aldrich) and 100 mM nicotinic acid. The synthesized ^32^P-NAADP was validated and purified with polyethyleneimine cellulose thin-layer chromatography developed in a solvent system of isobutyric acid−500 mM NH_4_OH (5:3 v/v). For the assay involving ^32^P-NAADP binding to purified hLsm12-His_E.coli_, the protein at 30 nM was incubated with various concentrations of non-radiolabeled NAADP or NADP for 30 min and then further incubated with 1 nM ^32^P-NAADP at room temperature for 1 h. The NAADP-protein complex was captured by Ni-NTA beads, washed three times with TBS, and eluted with 300 mM imidazole in TBS. For the assay involving ^32^P-NAADP binding to cell membranes, we followed a previously reported procedure^6^ with slight modifications. Cell membranes corresponding to 1 mg protein were incubated with 1 nM ^32^P-NAADP after preincubation with non-radiolabeled NAADP/NADP. Unbound ^32^P-NAADP was separated from the membrane through rapid vacuum filtration through GF/B filter paper. The radioactivity in the eluates or filter papers was counted with a liquid scintillation counter.

### Generation of Lsm12^Δ45-50^ mutant mice and preparation of MEFs

The founder lines of Lsm12 mutant mice were generated using the CRISPR/Cas9 method via pronuclear injection of a mixture of sgRNA 5’-CCAGAAUGUCCCUCUUCCAG-3’ (Synthego, unmodified) and Cas9 into embryonic stem cells derived from C57BL/6 mice at the MD Anderson Cancer Center Genetically Engineered Mouse Facility. Homozygous Lsm12^Δ45-50^ mutant mice and commercial C57BL/6 WT mice were used for preparation of MEFs. MEFs were isolated and cultured as previously described ^34^. Briefly, Embryos were isolated from E12.5 - E13.5 mouse embryos. After the head and most of the internal organs were removed, each embryo was minced and digested for 15 min, and then cultured at 37°C, 5% CO _2_ in freshly prepared MEF medium composed DMEM, 10% FBS, 2 mM L-glutamine and 100 U/ml penicillin-streptomycin. For Ca^2+^-imaging analysis, AAV1 particles (Addgene, Cat # 100836-AAV1) carrying CAG-driven GCaMP6f Ca^2+^ sensor was added in medium for 16 - 24 hours at 37°C. Cells were analyzed 24-48 hrs after a change of the medium to remove the virus. All animal experiments were carried out according to protocols and guidelines approved by the Institutional Animal Care and Use Committee of The University of Texas MD Anderson Cancer Center.

### Imaging analysis of NAADP-evoked Ca^2+^ release

Transfected cells were identified by fluorescence of the Ca^2+^ reporter GCaMP6f. Fluorescence was monitored with an Axio Observer A1 microscope equipped with an AxioCam MRm digital camera (Carl Zeiss) at a sampling frequency of 2 Hz. Cell injection was performed with a FemtoJet microinjector (Eppendorf). The pipette solution contained (mM): 110 KCl, 10 NaCl, and 20 Hepes (pH 7.2) supplemented with Dextran (10,000 MW)-Texas Red (0.3 mg/ml) and NAADP (100 nM) or vehicle. The bath was Hank’s Balanced Salt Solution (HBSS) that contained (mM): 137 NaCl, 5.4 KCl, 0.25 Na_2_HPO_4_, 0.44 KH_2_PO_4_, 1 MgSO_4_, 1 MgCl_2_, 10 glucose, and 10 HEPES (pH 7.4). When purified hLsm12-His_E.coli_ was used, it was injected at 100 ng/ml (∼ 4 μM) with NAADP in the same pipette solution. To minimize interference by contaminated Ca^2+^, the pipette solution was treated with Chelex 100 resin (Cat# C709, Sigma-Aldrich) immediately before use. Microinjection (0.5 s at 150 hPa) was made ∼30 s after pipette tip insertion into cells. Only cells that showed no response to mechanical puncture, i.e., no change in GCaMP6f fluorescence for ∼30 s, were chosen for pipette solution injection. Successful injection was verified by fluorescence of the co-injected Texas Red. The amount of injected solution was estimated to be much less than10% of the cell volume for each recorded cell. Elevation in intracellular Ca^2+^ concentration was reported by a change in fluorescence intensity ΔF/F_0_, calculated from NAADP microinjection-induced maximal changes (ΔF at the peak) in fluorescence divided by the baseline fluorescence (F_0_) immediately before microinjection.

### Patch-clamp recording of NAADP-induced TPC2 activation

Whole-cell patch-clamp recordings were performed on HEK293 cells with asymmetric Na^+^ (outside) /K^+^ (inside) solutions using a MultiClamp 700B amplifier (Axon Instruments) at room temperature. Bath solution contained 145 mM NaMeSO_3_, 5 mM NaCl, and 10 mM HEPES (pH 7.2). Pipette electrodes (3-5 MΩ) were filled with 145 mM KMeSO_3_, 5 mM KCl, and 10 mM HEPES (pH 7.2). The cells were visualized under an infrared differential interference contrast optics microscope (Zeiss). Currents were recorded by voltage ramps from -120 to +120 mV over 400 ms for every 2 s with a holding potential of 0 mV. After a whole cell recording configuration was achieved, an injection pipette was inserted into the cell and the baseline of the whole cell current was recorded. Microinjection of NAADP and purified Lsm12 protein was performed as above in imaging analysis of NAADP-evoked Ca^2+^ release. The NAADP-induced currents were obtained by subtraction of the baseline from NAADP injection-induced currents.

## ACKNOWLEDGMENTS

We thank Michael X. Zhu (The University of Texas Health Science Center at Houston) and Youxing Jiang (The University of Texas Southwestern Medical Center) for manuscript reading, discussion and comments. We thank Meggie Young and Kunal Shah for protein purification and all lab members for discussion. Mass spectrometry raw data were collected at the UT Southwestern Proteomic Core. The founder lines of mutant mice were generated at MD Anderson Cancer Center Genetically Engineered Mouse Facility. This work was supported by National Institutes of Health grants NS096296 (J.Y.) and GM130814 (J.Y.).

## AUTHOR CONTRIBUTIONS

J.Z. performed protein expression/characterization/isolation, proteomic analysis, and protein-ligand & protein-protein binding assays and initiated Ca^2+^-imaging and Lsm12 KO experiments. X.G. generated Lsm12-KO cells and mutant mice, performed immunofluorescence, performed microinjection, and collected Ca^2+^-imaging and electrophysiological data. J.Z. and J.Y. made protein expression constructs. J.Z. and X.G. generated and characterized Lsm12-KD cells. J.Y. conceived and supervised the project. J.Z., X.G., and J.Y. designed the research, analyzed data, and wrote the manuscript.

## COMPETING FINANCIAL INTERESTS

The authors declare that they have no competing interests.

## MATERIALS & CORRESPONDENCE

Correspondence and requests for materials should be addressed to J.Y.

## Notes

### Competing Interest Statement

The authors have declared no competing interest.

