## Supplemental Information for "Lsm12 is an NAADP receptor and a two-pore channel regulatory protein required for calcium mobilization from acidic organelles"

**Table S1. List of proteins as differential proteins between the test and control samples identified by quantitative mass spectrometric analysis from TPC1- and TPC2-expressing HEK293 cells.**

| **TPC1 interactome** | | | | | **TPC2 interactome** | | | | |
|---|---|---|---|---|---|---|---|---|---|
| **UniProt ID** | **Mass** | **Score[1]** | **# peptides[2]** | **H/L ratio[3]** | **UniProt ID** | **Mass** | **Score[1]** | **# peptides[2]** | **H/L ratio[3]** |
| TPC1 | 94,147 | 9619 | 164 | 10.1 | TPC2 | 85,243 | 6676 | 96 | 5.5 |
| SYFM | 52,357 | 159 | 2 | 15.4 | FARP1 | 118,633 | 230 | 5 | 55.1 |
| PRPS1 | 34,834 | 158 | 4 | 3.9 | TIM50 | 39,646 | 153 | 5 | 4.5 |
| MDHC | 36,426 | 128 | 2 | 4.6 | RT25 | 20,116 | 114 | 6 | 8.7 |
| **LSM12** | 21,701 | 126 | 2 | 1,586.3 | RAD50 | 153,892 | 105 | 2 | 3.9 |
| RT28 | 20,843 | 118 | 2 | 4.4 | TDRD3 | 73,185 | 98 | 2 | 9.5 |
| NEP1 | 26,720 | 114 | 2 | 4.1 | BMP2K | 129,172 | 96 | 2 | 3.2 |
| **FHAD1** | 161,904 | 106 | 6 | 8.3 | **C19L1** | 60,619 | 96 | 2 | 6.5 |
| TRI26 | 62,166 | 77 | 4 | 8.3 | **OBSCN** | 868,484 | 90 | 2 | 16.4 |
| PGRC1 | 21,671 | 76 | 2 | 18.8 | LIN7A | 25,997 | 84 | 2 | 278.7 |
| **SH3R1** | 93,129 | 74 | 8 | 29.1 | ATPO | 23,277 | 78 | 2 | 3.7 |
| TOP3A | 112,372 | 73 | 5 | 42.3 | C2D1A | 104,062 | 77 | 2 | 4.3 |
| S30BP | 33,870 | 72 | 2 | 2,084.3 | PEF1 | 30,381 | 77 | 1 | 75.4 |
| RT35 | 36,844 | 69 | 2 | 4.0 | 41 | 97,017 | 73 | 3 | 7.9 |
| **C19L1** | 60,619 | 66 | 2 | 3.2 | NOL12 | 24,663 | 67 | 3 | 30.4 |
| ERP44 | 46,971 | 65 | 4 | 10.1 | CCNL2 | 58,147 | 66 | 2 | 179.6 |
| **OBSCN** | 868,484 | 63 | 4 | 18,503.0 | **LSM12** | 21,701 | 66 | 4 | 6.8 |
| CGBP1 | 18,820 | 63 | 3 | 43.9 | MGME1 | 39,421 | 65 | 2 | 1,412.8 |
| CLP1 | 47,646 | 62 | 4 | 97.3 | ZO2 | 133,958 | 65 | 5 | 9.1 |
| IMDH1 | 55,406 | 58 | 4 | 13.0 | MIRO2 | 68,118 | 64 | 2 | 9.5 |
| PRS7 | 48,634 | 56 | 2 | 4.3 | **SH3R1** | 93,129 | 63 | 7 | 50.4 |
| PRG4 | 151,061 | 55 | 4 | 10,131.5 | CPNE1 | 59,059 | 62 | 1 | 9.7 |
| NB5R4 | 59,474 | 55 | 4 | 39.9 | SUCB2 | 46,511 | 61 | 4 | 44.5 |
| GAS8 | 56,356 | 52 | 2 | 16.1 | ZN700 | 86,232 | 61 | 4 | 4,612.0 |
| **RM20** | 17,443 | 51 | 1 | 5,180.0 | SSH1 | 115,511 | 56 | 3 | 9.6 |
| RM21 | 22,815 | 49 | 4 | 323.8 | PININ | 81,628 | 54 | 4 | 1,803.3 |
| **PRS6A** | 49,204 | 49 | 2 | 68.6 | DCAF7 | 38,926 | 53 | 1 | 22.2 |
| KINH | 109,685 | 46 | 2 | 6.4 | CENPP | 33,165 | 52 | 2 | 11.3 |
| RTN4 | 129,931 | 46 | 1 | 4.9 | HEXI2 | 32,419 | 51 | 1 | 269.6 |
| AT1A2 | 112,265 | 46 | 4 | 4.0 | **FHAD1** | 161,904 | 48 | 2 | 8.6 |
| NDUA5 | 13,459 | 45 | 1 | 38.0 | CAND1 | 136,376 | 45 | 1 | 7.0 |
| NOMO1 | 134,324 | 44 | 2 | 43.0 | LANC1 | 45,283 | 45 | 2 | 7.4 |
| ATP4A | 114,119 | 44 | 4 | 4.4 | WDR1 | 66,194 | 45 | 1 | 26.0 |
| LAMP2 | 44,961 | 43 | 1 | 3.0 | PUR2 | 107,767 | 43 | 2 | 78.6 |
| FIZ1 | 51,996 | 42 | 2 | 12,550.0 | CBX4 | 61,368 | 42 | 1 | 56.5 |
| **TCEA3** | 38,972 | 42 | 3 | 13.9 | GTF2I | 112,416 | 41 | 2 | 331.6 |
| ZN644 | 149,565 | 41 | 4 | 23.7 | LLPH | 15,225 | 41 | 1 | 3.3 |
| S10AB | 11,740 | 41 | 1 | 9.1 | **RM20** | 17,443 | 41 | 1 | 35.1 |
| FOLR3 | 27,638 | 40 | 1 | 42,100.0 | **PRS6A** | 49,204 | 40 | 1 | 235.8 |
| BIG2 | 202,038 | 40 | 2 | 6.2 | **TCEA3** | 38,972 | 40 | 2 | 658.0 |
| **NAADP interactome (TPC1-expressing cells)** | | | | | **NAADP interactome (TPC2-expressing cells)** | | | | |
| **UniProt ID** | **Mass** | **Score[1]** | **# peptides[2]** | **H/L ratio[3]** | **UniProt ID** | **Mass** | **Score[1]** | **# peptides[2]** | **H/L ratio[3]** |
| **CGL** | 44,508 | 439 | 9 | 5.6 | GSHR | 56,257 | 453 | 5 | 3.2 |

| | | | | |
|---|---|---|---|---|
| **KCC2D** | 56,369 | 363 | 6 | 3.8 |
| DHB4 | 79,686 | 229 | 1 | 3.8 |
| **BUB3** | 37,155 | 209 | 10 | 4.6 |
| **PTBP1** | 57,221 | 163 | 7 | 3.0 |
| SMRC2 | 132,879 | 145 | 5 | 3.5 |
| LRC47 | 63,473 | 139 | 2 | 3.3 |
| **ODPA** | 43,296 | 138 | 5 | 3.1 |
| LDHA | 36,689 | 129 | 2 | 12.0 |
| **LC7L3** | 51,466 | 128 | 2 | 7.8 |
| NCKP1 | 128,790 | 113 | 4 | 3.6 |
| NUFP2 | 76,121 | 109 | 5 | 4.8 |
| TBB3 | 50,433 | 109 | 2 | 12.2 |
| WDR48 | 76,210 | 109 | 2 | 3.1 |
| CAF1B | 61,493 | 104 | 3 | 3.2 |
| ILF2 | 43,062 | 103 | 2 | 3.6 |
| **FBX22** | 44,508 | 101 | 2 | 3.2 |
| LONM | 106,489 | 97 | 2 | 3.0 |
| **MCM5** | 82,286 | 97 | 6 | 3.4 |
| PDIA4 | 72,932 | 97 | 3 | 3.6 |
| HCFC1 | 208,732 | 92 | 2 | 3.2 |
| **DRG1** | 40,542 | 90 | 2 | 5.7 |
| CPSF7 | 52,050 | 87 | 3 | 3.6 |
| ACSL3 | 80,420 | 84 | 2 | 3.9 |
| LMF2 | 79,698 | 79 | 1 | 4.9 |
| TOE1 | 56,548 | 79 | 1 | 5.2 |
| CBR4 | 25,301 | 74 | 1 | 4.0 |
| **MCM3** | 90,981 | 74 | 2 | 4.1 |
| MPP6 | 61,117 | 74 | 5 | 3.7 |
| OSGEP | 36,427 | 74 | 1 | 5.9 |
| HPRT | 24,579 | 71 | 1 | 3.1 |
| KHDR1 | 48,227 | 70 | 4 | 3.9 |
| CPT1A | 88,368 | 69 | 1 | 214.3 |
| **SEPT2** | 41,487 | 68 | 2 | 3.0 |
| ANM1 | 42,462 | 67 | 2 | 3.8 |
| EMC1 | 111,759 | 67 | 3 | 3.9 |
| RPRD2 | 156,020 | 67 | 1 | 4.1 |
| **CARF** | 61,125 | 63 | 2 | 4.0 |
| CYTSA | 124,602 | 62 | 2 | 4.1 |
| AAAS | 59,574 | 61 | 1 | 4.4 |
| RBBP5 | 59,153 | 60 | 1 | 3.6 |
| **AATM** | 47,518 | 59 | 3 | 3.0 |
| RAI14 | 110,041 | 57 | 1 | 115.8 |
| SEPT7 | 50,680 | 56 | 2 | 4.9 |
| DMAP1 | 52,993 | 55 | 2 | 3.5 |
| AL1B1 | 57,206 | 54 | 2 | 3.9 |
| HS90A | 84,660 | 54 | 2 | 3.2 |
| **DDX47** | 50,647 | 53 | 1 | 3.5 |
| GNAS1 | 111,025 | 53 | 3 | 3.1 |
| RT27 | 47,611 | 52 | 1 | 3.2 |
| **CKAP2** | 76,987 | 51 | 2 | 3.6 |
| MFR1L | 31,957 | 51 | 1 | 3.1 |
| SYIM | 113,792 | 50 | 1 | 4.4 |
| CLAP2 | 141,064 | 48 | 1 | 10.7 |
| VDAC1 | 30,773 | 46 | 1 | 6.3 |
| GCDH | 48,127 | 43 | 1 | 4.1 |

| | | | | |
|---|---|---|---|---|
| DDX1 | 82,432 | 397 | 5 | 4.3 |
| PAIRB | 44,965 | 389 | 8 | 3.1 |
| HNRH2 | 49,264 | 330 | 6 | 3.1 |
| KIFC1 | 73,748 | 325 | 7 | 5.4 |
| **PTBP1** | 57,221 | 267 | 10 | 4.0 |
| **BUB3** | 37,155 | 266 | 14 | 4.7 |
| **ODPA** | 43,296 | 238 | 5 | 3.3 |
| KIF2C | 81,313 | 189 | 6 | 5.4 |
| **CGL** | 44,508 | 184 | 9 | 5.6 |
| HNRPL | 64,133 | 179 | 13 | 3.4 |
| IPYR2 | 37,920 | 170 | 3 | 3.9 |
| HNRPF | 45,672 | 168 | 3 | 3.2 |
| C1TM | 105,790 | 162 | 10 | 3.8 |
| HMGCL | 34,360 | 154 | 7 | 3.4 |
| **MCM3** | 90,981 | 134 | 3 | 3.2 |
| **FBX22** | 44,508 | 113 | 2 | 5.1 |
| <u>**LSM12**</u> | 21,701 | 110 | 2 | 3.3 |
| **SEPT2** | 41,487 | 109 | 1 | 3.3 |
| P5CR1 | 33,361 | 103 | 2 | 5.3 |
| WDR82 | 35,079 | 103 | 2 | 5.8 |
| MCCB | 61,333 | 93 | 2 | 4.1 |
| SYFB | 66,116 | 92 | 3 | 4.0 |
| **DRG1** | 40,542 | 91 | 2 | 6.0 |
| MIC60 | 83,678 | 90 | 2 | 3.2 |
| SRPRB | 29,702 | 88 | 3 | 3.2 |
| **MCM5** | 82,286 | 87 | 5 | 3.4 |
| GCP60 | 60,593 | 84 | 2 | 3.4 |
| **DDX47** | 50,647 | 82 | 1 | 3.9 |
| DLDH | 54,177 | 82 | 2 | 3.4 |
| FAD1 | 65,266 | 77 | 4 | 3.6 |
| STRAP | 38,438 | 75 | 1 | 3.5 |
| **AATM** | 47,518 | 72 | 3 | 4.2 |
| FND3A | 131,852 | 70 | 2 | 4.6 |
| **KCC2D** | 56,369 | 70 | 6 | 4.0 |
| TPX2 | 85,653 | 67 | 4 | 9.7 |
| MTDC | 37,895 | 66 | 2 | 27.6 |
| **CKAP2** | 76,987 | 65 | 3 | 4.8 |
| DDX6 | 54,417 | 64 | 3 | 3.3 |
| TCOF | 152,106 | 61 | 3 | 3.1 |
| KCC2A | 54,088 | 60 | 4 | 4.6 |
| **CARF** | 61,125 | 58 | 2 | 3.9 |
| RAB14 | 23,897 | 58 | 2 | 4.7 |
| MAVS | 56,528 | 56 | 1 | 4.5 |
| TCPQ | 59,621 | 56 | 1 | 3.2 |
| BT2A1 | 59,633 | 55 | 2 | 4.2 |
| **SPF45** | 44,962 | 55 | 3 | 5.2 |
| HXB9 | 28,059 | 54 | 1 | 5.1 |
| SRC | 59,835 | 54 | 1 | 5.7 |
| PDIA1 | 57,116 | 53 | 1 | 4.8 |
| TRM1L | 81,747 | 53 | 1 | 3.2 |
| **LC7L3** | 51,466 | 51 | 1 | 3.6 |
| EWS | 68,478 | 48 | 2 | 17.3 |
| ZC11A | 89,131 | 48 | 2 | 5.9 |
| HSP7E | 54,794 | 46 | 1 | 3.4 |
| SP1 | 80,693 | 46 | 1 | 7.6 |

| | | | | | | | | | |
|---|---|---|---|---|---|---|---|---|---|
| GLNA | 42,064 | 42 | 1 | 8.6 | SUCA | 36,250 | 46 | 1 | 4.9 |
| HNRPQ | 69,603 | 42 | 1 | 4.6 | SH3G1 | 41,490 | 45 | 1 | 3.0 |
| TBL1R | 55,595 | 42 | 1 | 7.2 | MRP | 19,529 | 42 | 1 | 3.1 |
| **LSM12** | 21,701 | 41 | 1 | 3.0 | ADAS | 72,912 | 40 | 2 | 3.7 |
| SCLY | 48,149 | 41 | 3 | 3.2 | NSDHL | 41,900 | 40 | 2 | 18,851.0 |
| **SPF45** | 44,962 | 41 | 3 | 4.3 | SPB6 | 42,622 | 40 | 2 | 3.3 |
| PARN | 73,451 | 40 | 1 | 5.4 | | | | | |

[1] Total mascot score for the protein.
[2] Number of matched H/L peptide pairs used for calculation of the protein's H/L ratio.
[3] Median of the heavy/light ratio of the protein.
Only proteins with H/L ratio of ≥3 and a mascot score ≥40 are included.
Protein names overlap between TPC1 and TPC2 interactomes and between TPC1- and TPC2-expressing NAADP interacteromes are highlighted in bold. Lsm12 is underlined as it is the only protein overlaps between TPC and NAADP interactomes.

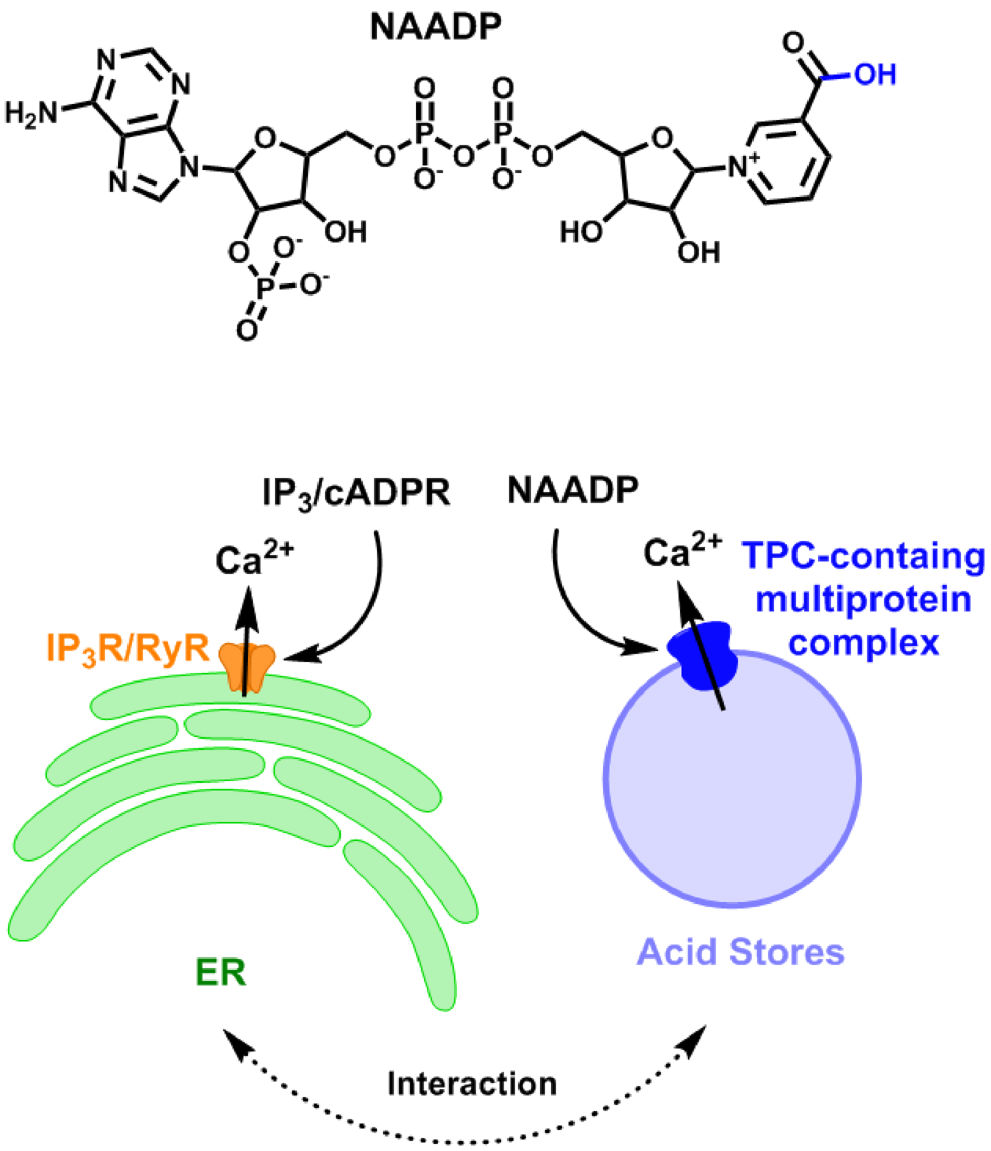


**Figure S1. Chemical structure of NAADP and $Ca^{2+}$-mobilizing second messenger pathways.** The -OH group, which differs from that (-$NH_2$) in NADP, is highlighted in blue in NAADP structure. A hypothetic TPC-containing multiprotein complex is proposed as the NAADP signaling complex responsible for NAADP-evoked $Ca^{2+}$ release from acid stores.

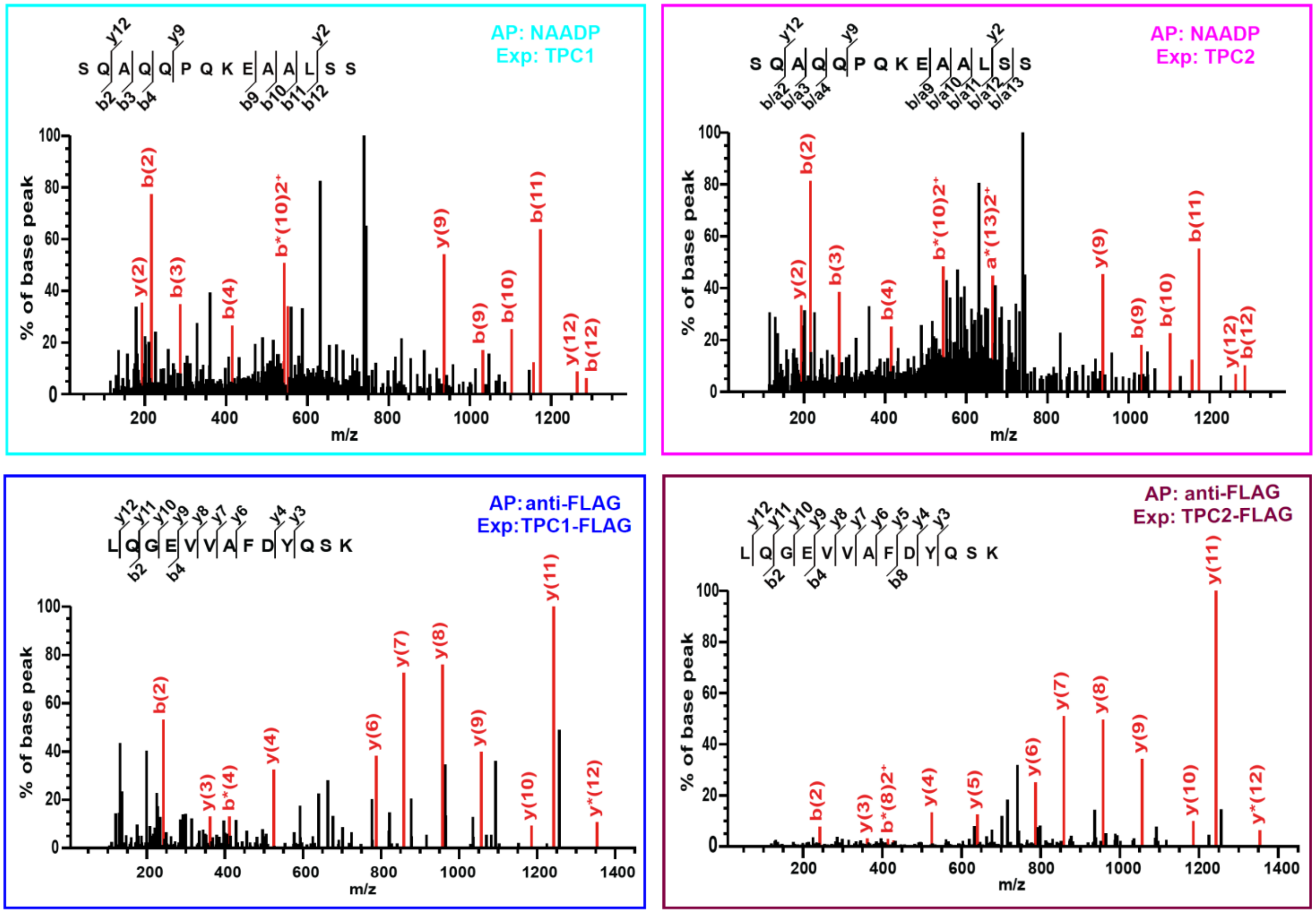


**Figure S2. MS/MS spectra of Lsm12 peptides.** MS/MS spectra of the peptide SQAQQPQKEAALS[194] and LQGEVVAFDYQSK[37] corresponding to those in Figure 1d. AP, affinity precipitation. Exp, expression.

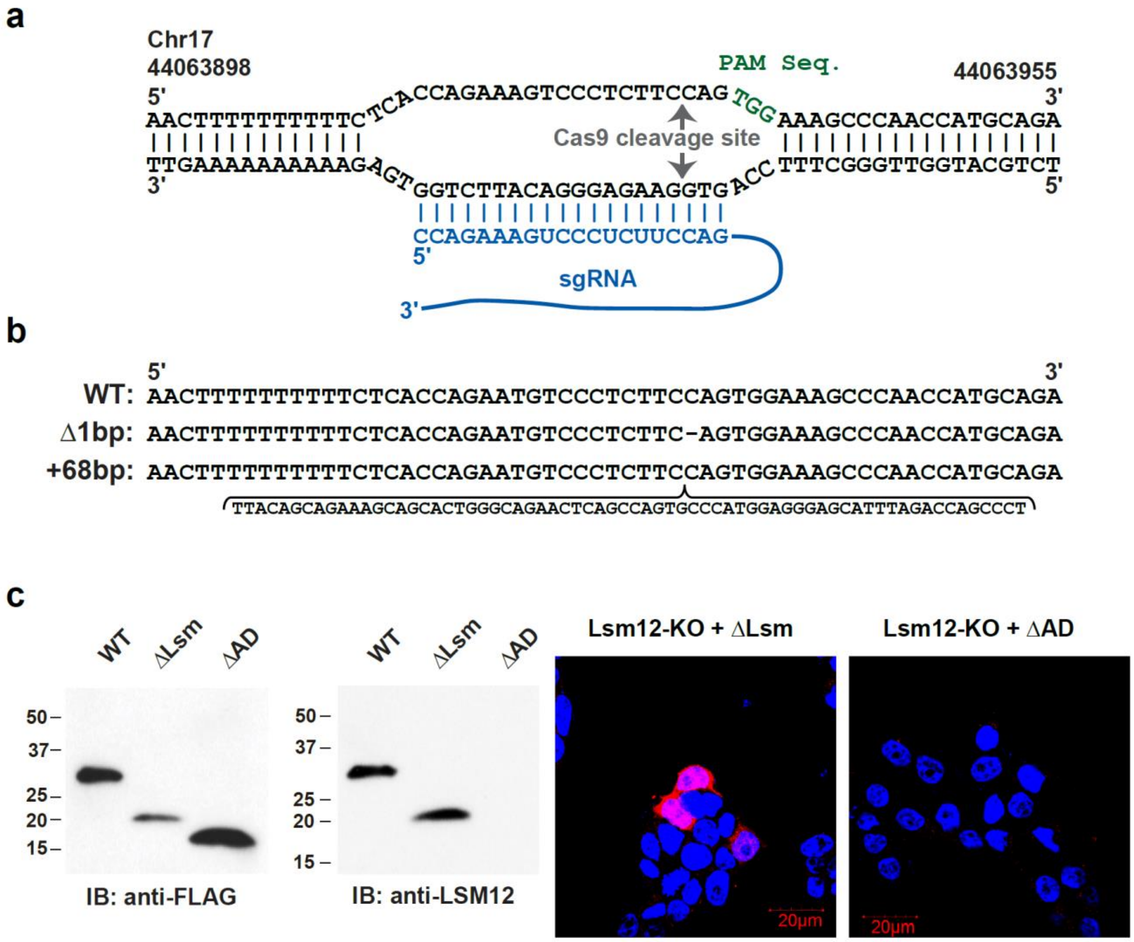


**Figure S3. Generation and characterization of the Lsm12-KO line of HEK293 cells. a**, Depiction of sgRNA-genomic DNA complex showing the targeted site of genomic editing in Lsm12 gene. **b**, Nucleotide sequences of Lsm12 genomic DNA in the sgRNA targeted region in HEK293 WT and Lsm12-KO cells. **c**, Immunoblot and immunofluorescence of Lsm12 by an anti-Lsm12 antibody of Lsm12-KO cells transiently expressing exogenous FLAG-tagged Lsm12-WT, -ΔLsm and -ΔAD mutants. Same amount of total protein of cell lysates was loaded for each lane in SDS-PAGE.

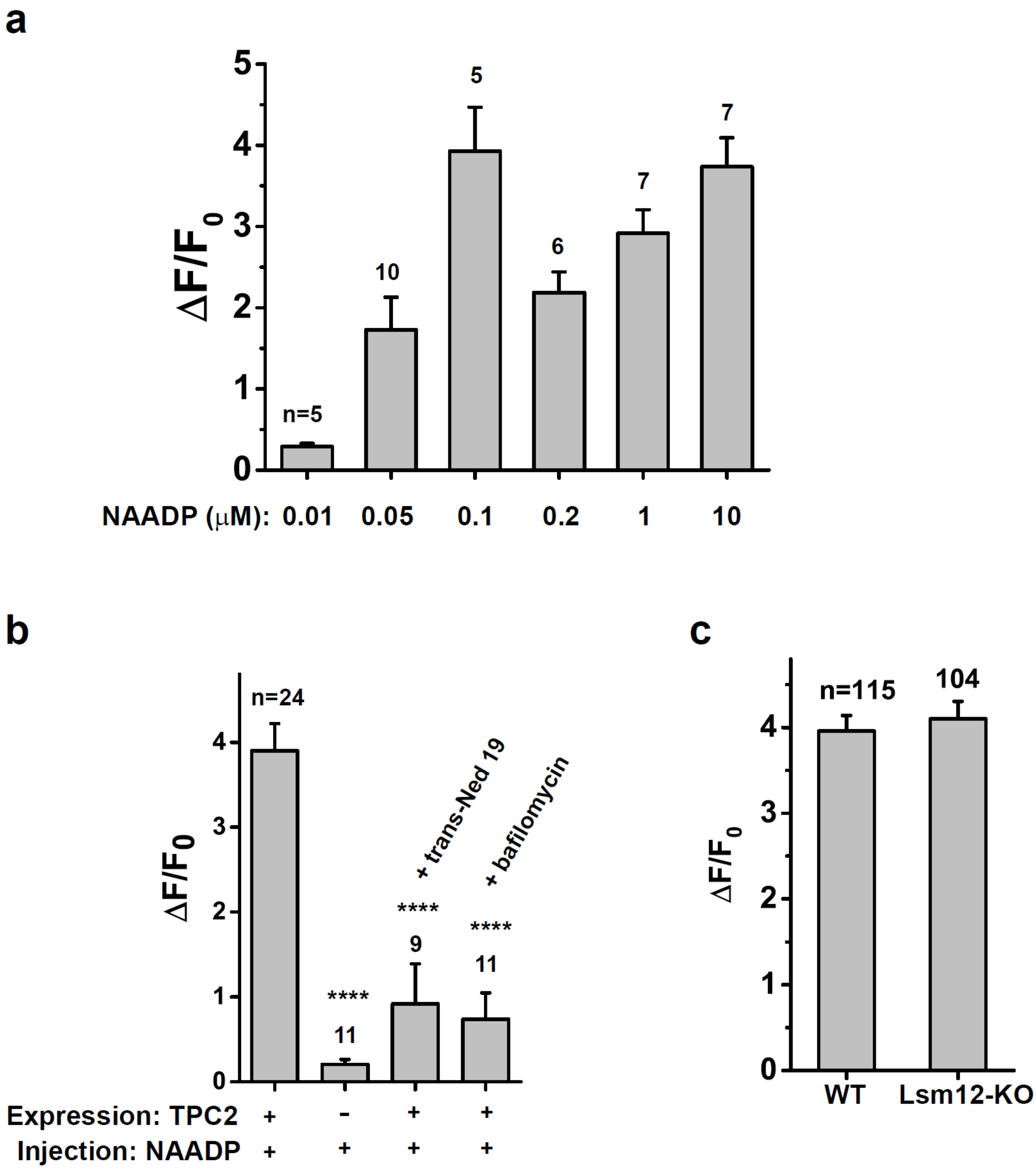


**Figure S4. NAADP-evoked $Ca^{2+}$ changes in HEK293 WT cells under different microinjection or cell treatment conditions. a**, The effect of different NAADP concentrations in injection pipette solution on microinjection-induced change in the $Ca^{2+}$ indicator fluorescence. **b**, NAADP (100 nM in pipette solution) microinjection-evoked change in the $Ca^{2+}$ indicator fluorescence under different conditions. The cells were treated with bafilomycin A1 at 1 µM or trans-Ned 19 at 10 µM for 45- 60 min in cell growth medium immediately before $Ca^{2+}$ imaging. **c**, Extracellular ATP treatment–induced changes in intracellular $Ca^{2+}$ were similar in WT and Lsm12-KO cells. ATP was manually added to the bath solution at a final concentration of 50 µM. The number of cells analyzed are indicated.

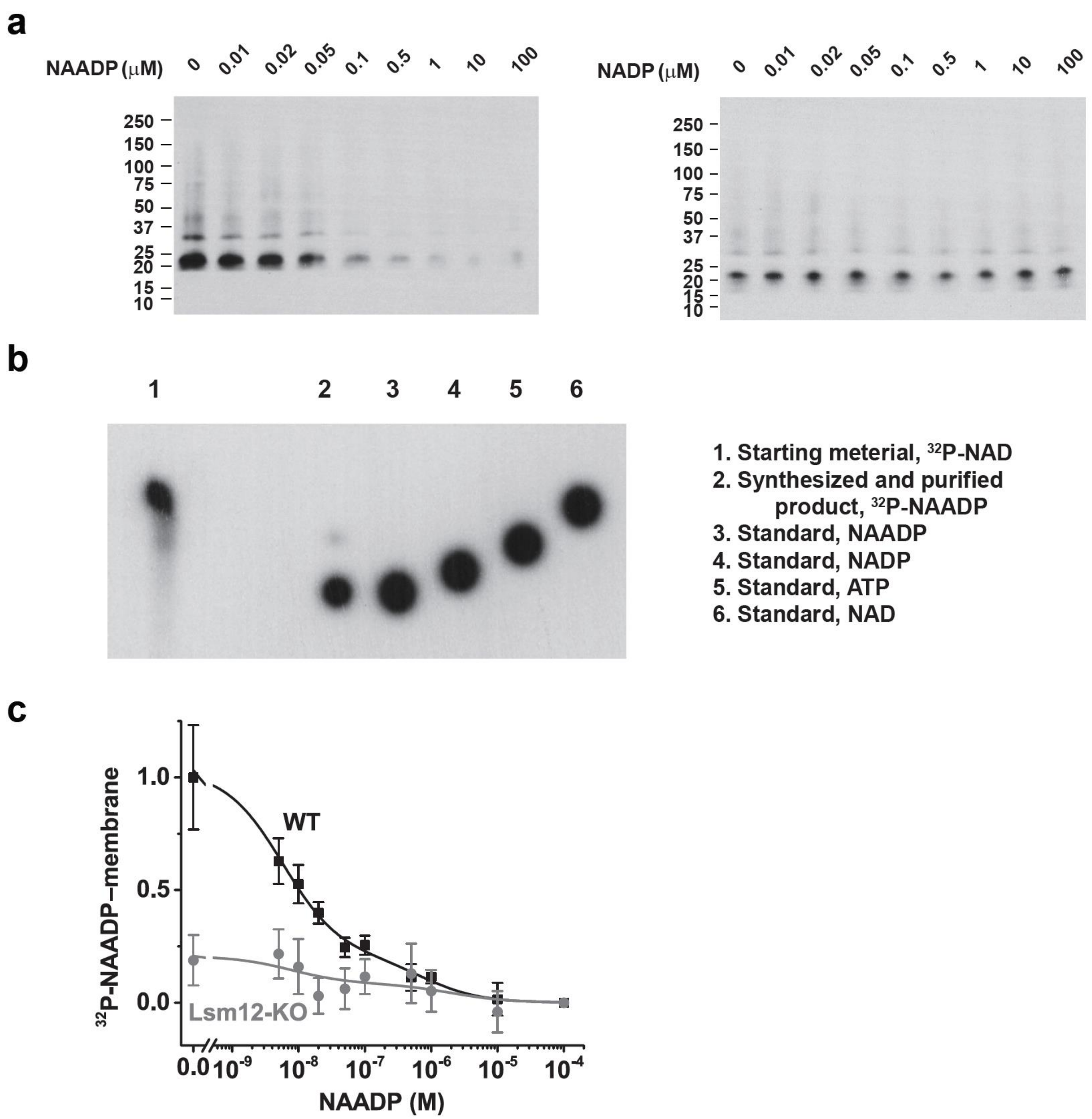


**Figure S5. Competition binding assay of NAADP binding to hLsm12-His$_{E.coli}$ and TPC2-expressing HEK293 cell membranes. a,** Immunoblot of hLsm12-His$_{E.coli}$ pulled down by immobilized NAADP in the absence or presence of various concentrations of free NAADP and NADP. **b**, Thin-layer chromatography (TLC) of synthesized and purified $^{32}$P-NAADP. The image was obtained by autoradiography. The standard NAADP, NADP, ATP, and NAD were visualized under a UV lamp on TLC plate first and then spotted with $^{32}$P. **c**, Competition radioligand binding assay of the association between $^{32}$P-NAADP and TPC2-expressing cell membranes in the presence of various concentrations of non-radiolabled NAADP.

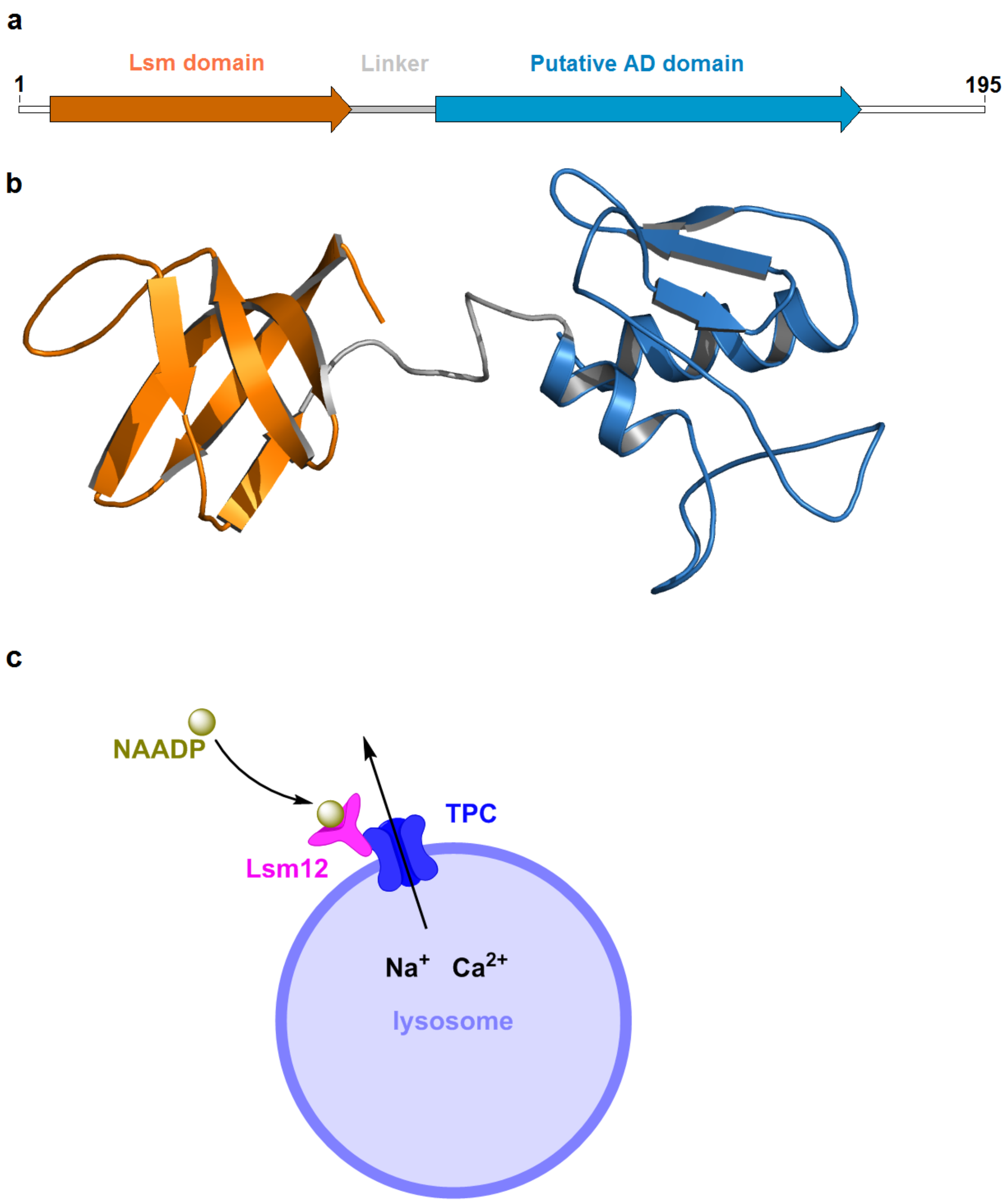


**Figure S6. Features of Lsm12. a,** Putative domain assignment for Lsm12. **b**, A predicted 3-D structural model of human Lsm12 drawn with a previously created model from the SWISS MODEL Repository (https://swissmodel.expasy.org/repository/uniprot/Q3MHD2) [1], modeled with a structural template [2] of an archaea atypical Lsm protein called SmAP3. **c**, Proposed model of Lsm12 in NAADP signaling.

1 Bienert, S. *et al.* The SWISS-MODEL Repository-new features and functionality. *Nucleic Acids Research* **45**, D313-D319, doi:10.1093/nar/gkw1132 (2017).
2 Mura, C., Phillips, M., Kozhukhovsky, A. & Eisenberg, D. Structure and assembly of an augmented Sm-like archaeal protein 14-mer. *Proceedings of the National Academy of Sciences of the United States of America* **100**, 4539-4544, doi:10.1073/pnas.0538042100 (2003).
